# Connecting mitochondrial metabolism and mitotic fidelity to control vulnerability of high grade serous ovarian cancer patients to taxane-based chemotherapy

**DOI:** 10.64898/2026.01.07.698258

**Authors:** Hadia Moindjie, Morgane Morin, Evanthia Pangou, Diego Chianese, Cynthia Seiler, Sylvie Rodrigues-Ferreira, Maria M. Haykal, Christine Péchoux, Florent Dingli, Damarys Loew, Yann Kieffer, Geraldine Gentric, Patricia Pautier, Philippe Dessen, Rafael José Argüello, Paolo Pinton, Fatima Mechta-Grigoriou, Catherine Brenner, Massimo Bonora, Clara Nahmias

**Author notes:** Equal contribution (Morgane Morin, Evanthia Pangou, Diego Chianese). To whom correspondence may be sent (Clara Nahmias). Present address Université Paris-Saclay, UVSQ, Inserm, IMPROVE, 78000 Versailles, France.

## Abstract

High-grade serous ovarian carcinoma (HGSOC), which accounts for approximately 75% of ovarian cancer cases, is associated with poor clinical outcome. Although most patients initially achieve a complete response to conventional chemotherapy, HGSOC almost invariably develops chemoresistance. There is therefore an urgent need to identify predictive biomarkers of treatment response. Here, through integrative analyses of molecular and clinical data from HGSOC patient cohorts, we identify syntabulin (SYBU), a microtubule-associated protein originally described as a regulator of mitochondrial transport along neuronal microtubules, as a critical determinant of chemosensitivity in HGSOC. Low SYBU expression in tumors correlates with higher tumor grade and increased aggressiveness, yet paradoxically with enhanced sensitivity to chemotherapy. SYBU-deficient cancer cells display impaired oxidative phosphorylation and a metabolic shift toward glycolysis characteristic of the Warburg effect, together with mitotic defects such as chromosome lagging that promote aneuploidy. Mechanistically, syntabulin forms a complex with the mitochondrial outer membrane porin VDAC1 and the inner membrane protein MIC60, a major regulator of mitochondrial cristae organization. Functionally, the syntabulin-MIC60 axis controls cristae architecture and mitotic fidelity, thereby connecting mitochondrial metabolism to cell division. These findings highlight new therapeutic vulnerabilities to overcome chemoresistance in ovarian cancer.

**SIGNIFICANT STATEMENT:** Ovarian cancer remains the deadliest gynecologic malignancy, largely due to the systematic emergence of resistance to chemotherapy. Identifying molecular mechanisms involved in response to treatment is therefore a major clinical challenge. Here, we uncover an unexpected role for the mitochondrial protein syntabulin in regulating chemotherapy sensitivity in high-grade serous ovarian cancer. We demonstrate that syntabulin coordinates cancer cell mitotic progression with mitochondrial structure and metabolism through interactions with cristae-shaping proteins. These findings reveal a previously unrecognized link between mitotic regulation and mitochondrial architecture, and identify syntabulin as a potential therapeutic target in ovarian cancer to induce vulnerability to taxane-based chemotherapy.

## INTRODUCTION

Ovarian cancer is the fourth cause of cancer death in women worldwide and remains the deadliest gynecologic malignancy. In 2020, nearly 310,000 women were diagnosed with ovarian cancer, and approximately 200,000 died from the disease. High-grade serous ovarian cancer (HGSOC), the most common form of ovarian cancer representing 75% of cases, has a 5-year overall survival of less than 50% [1,2]. Standard treatment for HGSOC relies on neoadjuvant platinum–taxane chemotherapy followed by debulking surgery [3–5]. However, this treatment induces undesirable toxicities and the majority of patients eventually develop initial or acquired resistance to chemotherapy, leading to disease progression and death [6–8]. In the era of precision medicine, the identification of new predictive biomarkers of HGSOC chemoresistance is urgently needed in order to stratify the patients who may benefit from the treatment and provide a close clinical follow-up for those who remain resistant [9–12].

Conventional taxane-based chemotherapy - including paclitaxel (PTX) and docetaxel - targets the microtubule cytoskeleton by stabilizing microtubules, thereby disrupting mitotic spindle dynamics, leading to mitotic arrest and ultimately triggering cell death. The assembly, dynamics and functions of the microtubule network are orchestrated by a large number of microtubule-regulatory (MT-reg) proteins that control cellular homeostasis [13,14]. We hypothesized that any alteration in the expression levels of MT-reg genes in HGSOC tumors may trigger remodeling of the microtubule cytoskeleton and have important effects on tumor sensitivity to taxanes. In line with this hypothesis, dysregulation of genes encoding microtubule-associated proteins in chemoresistant breast cancer patients has previously been reported [14–16].

In this study, we analyzed the expression levels of 410 genes encoding MT-reg proteins to evaluate their predictive value in two independent cohorts of HGSOC patients treated with neoadjuvant taxane-based chemotherapy. *SYBU*, the most differentially expressed gene between sensitive and resistant tumors in both cohorts of patients, was selected for further studies. *SYBU* encodes syntabulin, a mitochondrial protein originally characterized in neurons where it is involved in anterograde transport of mitochondria along microtubules [17–19]. In breast cancer, syntabulin also controls mitochondrial transport along microtubules to regulate cell migration and cancer metastasis [20]. To date, the contribution of *SYBU* to ovarian cancer biology and chemoresistance has remained unexplored. Here, we demonstrate that low levels of *SYBU* in HGSOC patients are associated with increased sensitivity to chemotherapy. At the cellular level, *SYBU* depletion induces mitotic defects and reprograms mitochondrial metabolism, which constitute two major vulnerabilities to taxane treatment. At the molecular level, we show that syntabulin associates with mitochondrial outer and inner membrane proteins VDAC1 and MIC60. Together our data uncover a syntabulin-MIC60 axis that connects mitochondrial architecture to mitotic fidelity, with direct implications for taxane-based chemotherapy response in HGSOC.

## RESULTS

### Low *SYBU* levels are associated with sensitivity to chemotherapy in HGSOC patients

To investigate the potential value of MT-reg genes as predictive biomarkers of ovarian cancer chemoresistance, we focused on a series of 410 genes (Suppl Table TS1) encoding known regulators of microtubule organization or function. Expression levels of these genes were analyzed in two independent cohorts of ovarian cancer patients from Australia (AOCS cohort) and France (CURIE cohort). Patients were selected based on high grade and serous histological subtype (Suppl Table TS2) and classified as sensitive (S) or resistant (R) to conventional chemotherapy (Suppl Fig. S1A) as described in the Methods. A total of 25 genes from the AOCS cohort and 20 genes from the CURIE cohort were found differentially expressed between groups of (S) and (R) patients (Suppl Table TS3, Fig.1A). Three genes (*MICAL1*, *KDM6A* and *SYBU*) were significantly differential in both cohorts. Among them, *SYBU* (also known as *GOLSYN*) was the most strongly differentially regulated gene as shown in volcanoplots (Fig. 1B) with a fold change of 0.61 and 0.23 in (S) versus (R) tumors of the AOCS and CURIE cohorts, respectively (Suppl Table TS3).

**Figure 1.**
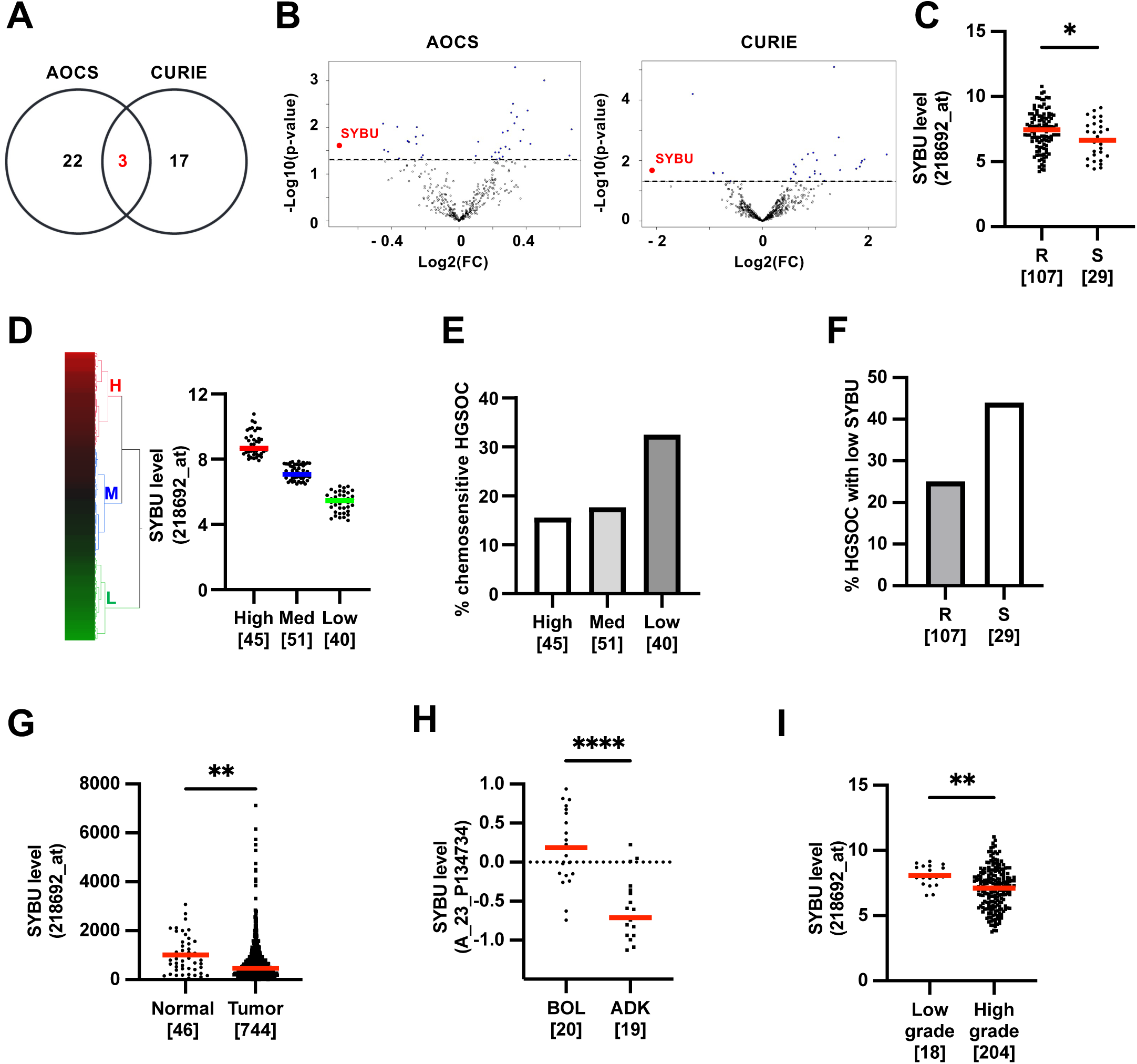
Identification of *SYBU* as a down-regulated gene in HGSOC patients. A. Venn diagram showing the number of differentially expressed genes (FC ≥ 1.2) between sensitive and resistant HGSOC in AOCS and CURIE cohorts. B. Volcanoplot representation of differentially expressed genes in AOCS (left) and CURIE (right) cohorts. Data are plotted as Log2 fold-change (FC) versus minus Log10 of the adjusted p-value. The vertical dashed line indicates the significance threshold (p-value = 0.05). Blue points above this line represent significantly deregulated genes (p < 0.05). Note the prominent position of the *SYBU* gene (red dot). C. Scattered dot plot of *SYBU* probeset (218692_at) intensities in tumors from patients of the AOCS cohort classified according to resistance (R) or sensitivity (S) to chemotherapy. Median values are highlighted in red. Number of tumors in each group is indicated in brackets. D. Heatmap and hierarchical clustering of 136 HGSOC samples based on the intensities of *SYBU* probeset (218692_at) in the AOCS cohort. The heatmap illustrates relative expression profiles of SYBU (column) for each tumor sample (line) in a continuous color scale from low (green) to high (red) expression. A dendrogram of the 3 selected tumor clusters (H for High, M for Medium and L for Low) is shown on the right side. Scattered dot plot of SYBU expression in each of the 3 selected clusters based on the dendrogram is shown in right panel. Number of tumors in each group is indicated in brackets. E. Percentage of tumors that are sensitive to chemotherapy among selected clusters of HGSOC expressing High, Medium or Low *SYBU* levels according to heatmap clustering (D). Number of tumors in each group is indicated in brackets. F. Percentage of tumors with low *SYBU* levels in resistant (R) and sensitive (S) HGSOC groups in the AOCS cohort. Number of tumors in each group is indicated in brackets. G. Scattered dot plot of the *SYBU* Affymetrix probeset (218692_at) intensities in normal and tumoral ovarian tissues according to TNMplot.com. Median values are highlighted in red. Number of tumors in each group is indicated in brackets. H. Scattered dot plot of *SYBU* Agilent probeset (A_23_P134734) intensities in borderline (BOL) and adenocarcinoma (ADK) ovarian tissues in the GR cohort. Median values are highlighted in red. Number of tumors in each group is indicated in brackets. I. Scattered dot plot of *SYBU* Affymetrix probeset (218692_at) intensities in low-grade and high-grade serous ovarian tumors in the AOCS cohort. Median values are highlighted in red. Number of tumors in each group is indicated in brackets. *P<0.05; **P<0.01; ****P<0.001.

In both cohorts of patients, *SYBU* Affymetrix probeset intensities were significantly higher in the (R) versus (S) group of HGSOC tumors (Fig. 1C and Suppl Fig. S1B). Heat map hierarchical clustering was used to classify samples of the AOCS cohort into three groups according to high (H), medium (M) and low (L) levels of *SYBU* expression (Fig. 1D) and their clinical data were examined. As shown in Fig. 1E, 32% of low-*SYBU* tumors were sensitive to chemotherapy, compared to 16% and 18% of high- and medium-*SYBU* expressing tumors, respectively. The percentage of tumors with low *SYBU* levels reached 25% in the (R) group, compared with 44% in the (S) group of patients (Fig. 1F). Similar results, showing marked association between low levels of *SYBU* and tumor sensitivity to chemotherapy, were obtained with the CURIE cohort (Suppl Fig. S1C-S1E). These results indicate that low-*SYBU* tumors respond better to treatment and point to the *SYBU* gene as a potential predictive biomarker of HGSOC sensitivity to chemotherapy.

To broaden our understanding of SYBU gene regulation in cancer, we turned to the publicly available cancer TNMplot database [21]. We found that *SYBU* probeset intensities were significantly lower in ovarian tumors compared with healthy ovarian tissue (Fig. 1G).

Among ovarian tumors, *SYBU* levels were significantly lower in malignant adenocarcinoma (ADK) compared with benign borderline (BOL) tumors (Fig. 1H, Suppl Table TS4). Histological subtypes of ovarian cancers with most aggressive properties and worse prognosis, such as Clear Cell and Serous Epithelial Ovarian cancers, were found to exhibit the lowest *SYBU* levels (Suppl Fig. S1F). Finally, serous epithelial ovarian tumors of high-grade were found to express lower levels of *SYBU* compared to those of low grade (Fig. 1I). Together, these results indicate that low *SYBU* levels are associated with increased ovarian cancer malignancy.

### SYBU depletion sensitizes cancer cells to paclitaxel

Taxanes are microtubule-targeting agents which, when used at nanomolar concentrations, induce several mitotic defects including centrosome amplification, multipolar spindle formation and chromosome missegregation [14,22–24]. Given that syntabulin, the S*YBU* gene product, is a microtubule-associated mitochondrial protein [18], we hypothesized that *SYBU* depletion may influence cancer cell sensitivity to taxanes by perturbing mitotic progression. To test this, HeLa cells - widely used as a model system to study cell division - were transfected with specific siRNAs (suppl Fig. S2A) and subsequently treated with low doses of paclitaxel (PTX) prior to analyzing their mitotic characteristics and cell fate. As shown in Fig. 2A and 2B, in untreated cells, *SYBU* silencing impaired chromosome alignment in metaphase and induced lagging chromosomes in anaphase and telophase. *SYBU* depletion also significantly increased the percentage of cells with supernumerary centrosomes (Fig. 2C), a phenotype associated with PTX treatment [14]. SYBU depletion thus mimics the deleterious effects of taxanes on mitotic features.

**Figure 2.**
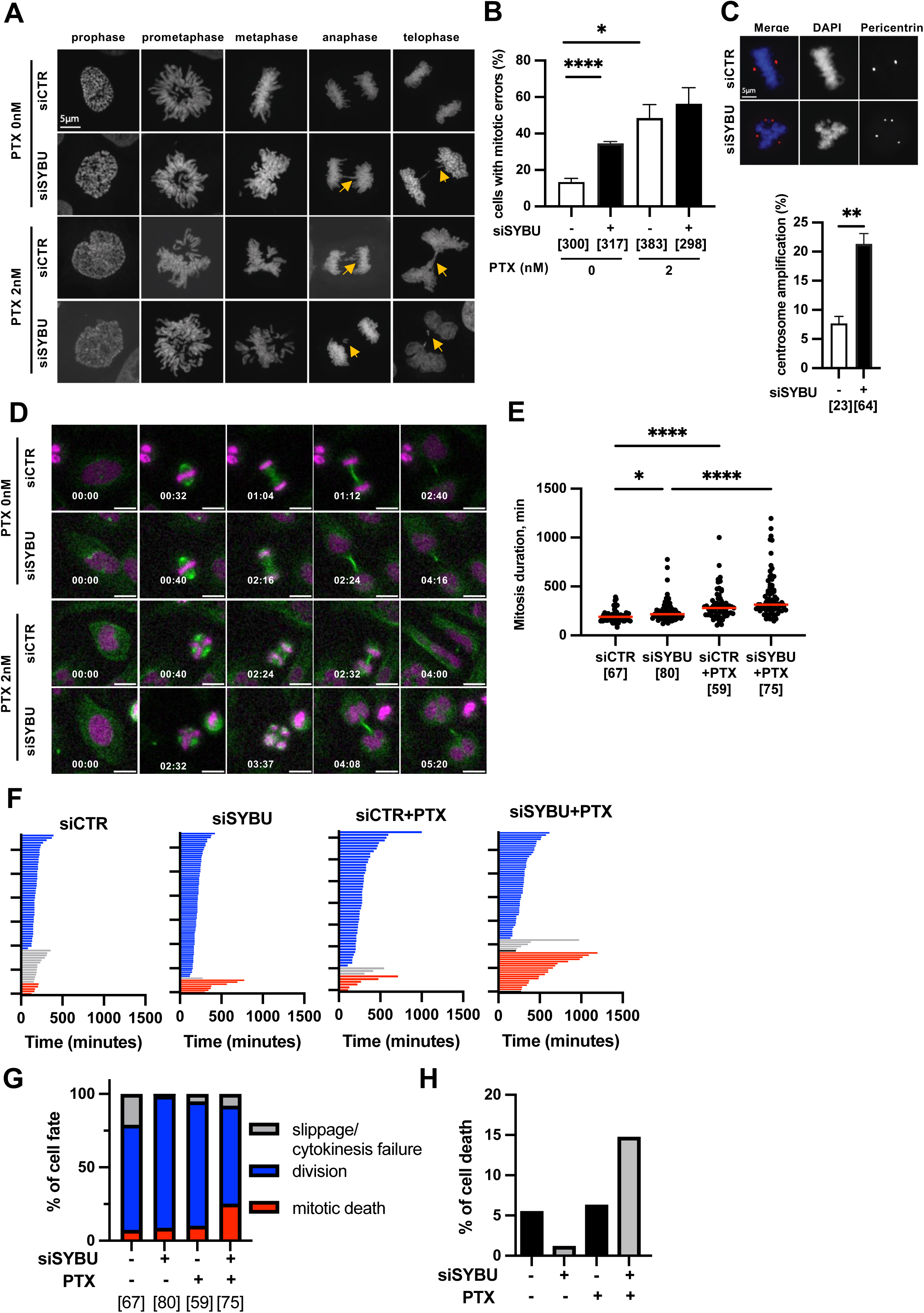
*SYBU* depletion induces mitotic defects and sensitizes HGSOC cells to PTX. A. Immunofluorescence photographs of HeLa cells expressing (siCTR) or not (siSYBU) endogenous *SYBU* and treated or not with 2 nM PTX for 10 hrs prior staining with DAPI. Arrows indicate lagging chromosomes in anaphase and telophase. Magnification, 63x. Scale bar = 5 μm B. Percentage of HeLa cells with mitotic errors shown in (A). C. Immunofluorescence photographs of HeLa cells expressing (siCTR) or not (siSYBU) endogenous syntabulin, stained with anti-pericentrin antibody (red) and DAPI (blue) to reveal centrosome and DNA respectively (top panel). Percentage of mitotic HeLa cells containing more than 2 centrosomes (bottom panel). D. Time-lapse fluorescent imaging of HeLa mCherry-H2B cells expressing (siCTR) or not (siSYBU) endogenous *SYBU*, stained with siR-tubulin and treated or not with 2nM PTX. Time is indicated on the bottom right as hrs:min. Microtubules (siR-tubulin) are shown in green and DNA (mCherry-H2B) in pink. Magnification, 20x. E. Scattered dot plot showing mitosis duration. F. Cell fate profiles of control (siCTR) or *SYBU*-depleted (siSYBU) HeLa mCherry-H2B cells on the first mitosis after treatment or not with PTX (2 nM). G. Percentage of cell fate after the first mitosis shown in (F). H. Percentage of cell death in interphase. (B-I). The number of cells analyzed is indicated in brackets. \**P* < 0.05; \*\**P* < 0.01; \*\*\*\**P* < 0.0001.

Cell fate was then evaluated by time-lapse videomicroscopy in HeLa-mCherry-H2B cells expressing fluorescent histone H2B (Fig. 2D). *SYBU* depletion slightly prolonged the time spent in mitosis and this was significantly increased in the presence of PTX (Fig. 2E, 2F). *SYBU* depletion markedly increased the percentage of mitotic death in response to PTX (Fig. 2F, 2G) and sensitized to PTX-induced cell death in interphase (Fig. 2H). Together, these results indicate that decreasing *SYBU* levels impairs mitotic progression and synergizes with taxane treatment to increase cell death, thereby rendering cancer cells more vulnerable to taxane-based chemotherapy.

### Loss of *SYBU* promotes metabolic vulnerability in HGSOC cells

Mitochondrial metabolism has been shown to govern the fidelity of mitotic progression [25–27], leading us to investigate whether syntabulin, which is anchored in the mitochondrial outer membrane, may contribute to mitochondrial metabolism. To address this question, we used COV318 and OVCAR8 cell lines derived from high grade epithelial serous ovarian cancer [28,29] that express detectable levels of *SYBU* (Suppl Fig.S2B).

Mass spectrometry-based metabolomic profiling of COV318 cells revealed that *SYBU* depletion using siRNA (Suppl Fig. S2C, S2D) induces significant metabolic reprogramming that affects mitochondrial metabolism, including fatty acid and butanoate metabolism, Coenzyme A biosynthesis and TCA cycle (Fig. 3A left panel); suppl Fig. S3A left panel). Among 56 metabolites analyzed, a total of 15 were significantly differential following *SYBU* depletion and/or PTX treatment (Fig. 3B, 3C; suppl Fig. S3B; Table TS5). Notably, Acetyl-CoA was found markedly accumulated in *SYBU*-deficient cells (Fig. 3C, 3D), suggesting potential alteration in mitochondrial metabolism. Metabolic rewiring was further exacerbated upon PTX treatment of *SYBU*-silenced cells (Fig.3A). In these cells, metabolites differentially regulated were enriched in pathways associated with the Warburg effect [30], highlighting a prominent role for syntabulin in the control of energy metabolism (Fig.3A right panel; suppl Fig.S3A right panel, Suppl Fig. S3B). Collectively, these data suggest that *SYBU* depletion establishes a metabolically vulnerable state characterized by altered carbon routing and mitochondrial function, revealing heightened metabolic plasticity of *SYBU*-deficient cells upon chemotherapeutic stress.

**Figure 3.**
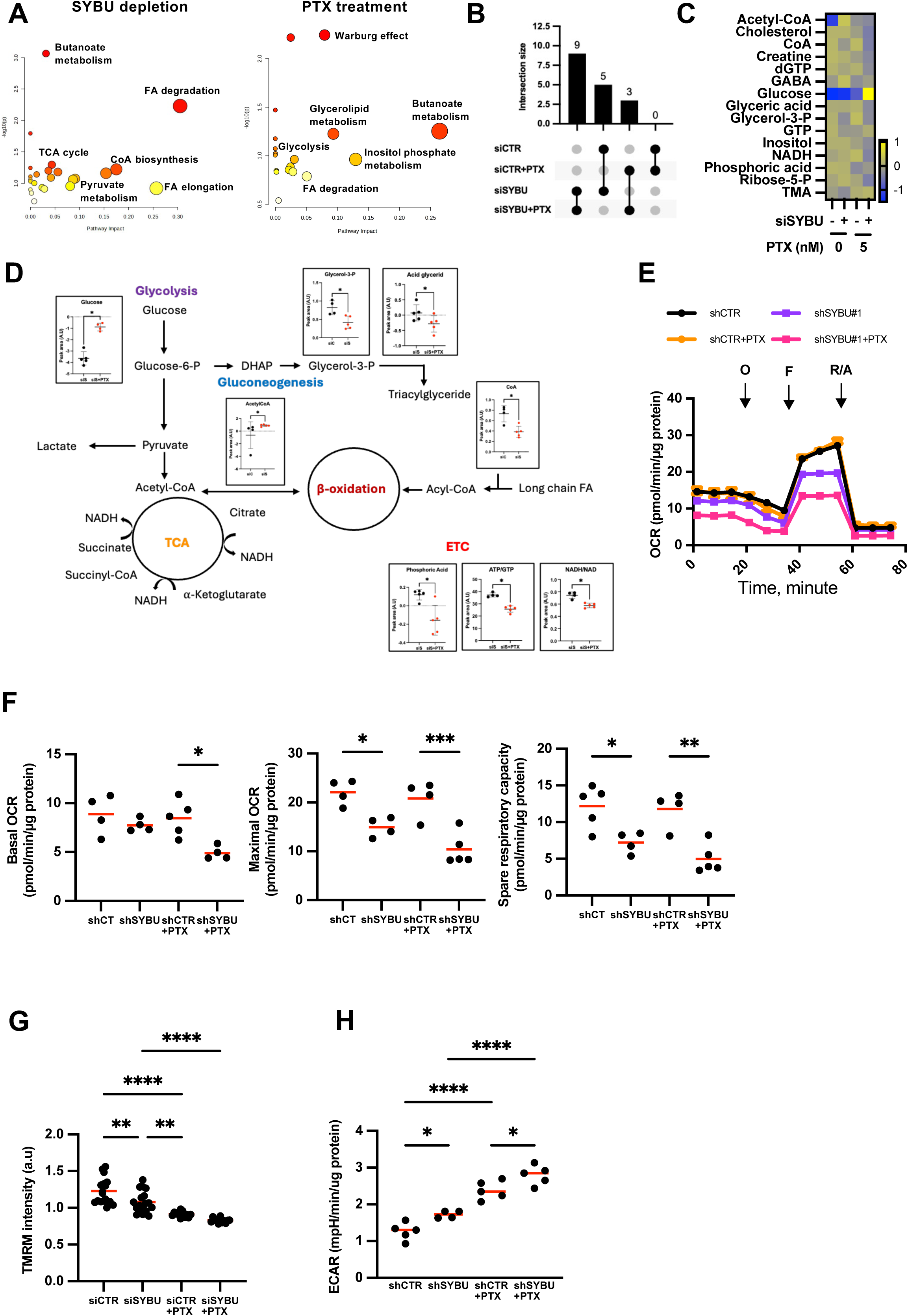
*SYBU* depletion impairs oxidative phosphorylation and promotes a metabolic shift toward glycolysis. A. Identification of metabolic pathways altered in COV318 cells following *SYBU* silencing (left panel) and in *SYBU*-silenced cells following PTX (5 nM) treatment for 72hrs (right panel) using MetaboAnalyst. B. UpSet plot illustrating shared significantly differential metabolites in cells expressing SYBU (siCTR) or not (siSYBU) and treated or not with 5nM PTX for 72 hrs. C. Heatmap of significant metabolites identified from metabolomic analysis. D. Peak area value (A.U, arbitrary units) and associated metabolic pathways for metabolic intermediates identified in metabolomic analysis. E. Representative curves of oxygen consumption rates (OCR) in COV318 cells permanently depleted (shSYBU#1) or not (shCTR) for *SYBU* using shRNA and treated or not with 5nM PTX for 72hrs. O: oligomycin, F: FCCP, and R/A: rotenone/antimycin were sequentially injected to assess mitochondrial respiratory states. 3–5 technical replicates per group. F. Quantification of basal respiration (left), maximal respiration (middle) and spare respiratory capacity (right panel) in COV318 cells from (E). G. Quantification of relative TMRM fluorescence intensity (a.u, arbitrary units) in COV318 cells expressing (siCTR) or not (siSYBU) *SYBU* and treated or not with 5nM PTX for 72 hrs. H. Quantification of extracellular acidification rate (ECAR) in COV318 cells permanently depleted (shSYBU) or not (shCTR) for *SYBU* and treated or not with 5nM PTX for 72 hrs. \**P* < 0.05; \*\**P* < 0.01; \*\*\**P* < 0.001; \*\*\*\**P* < 0.0001.

To further explore and confirm the functional consequences of SYBU depletion on mitochondrial capacities, we measured oxygen consumption rates (OCR) in ovarian cancer cells by seahorse analysis. In both COV318 and OVCAR8 cell lines, *SYBU*-depletion led to lower OCR compared with control (Fig. 3E, 3F; suppl Fig. S2C-S2F, S4A-S4F). Exposure to low doses of PTX in *SYBU*-depleted cells further reduced mitochondrial respiration (Fig. 3E, 3F). Of note, reduced mitochondrial respiration in SYBU depleted cells was not associated with a decrease of mitochondrial mass monitored both by MitoTracker staining and quantification of mitochondrial DNA (Suppl Fig. S4G, S4H) indicating that *SYBU* controls mitochondrial function but not biogenesis. Moreover, no significant variation in OXPHOS-complex proteins was detected upon *SYBU* depletion and treatment with PTX (suppl Fig. S4I). We then measured mitochondrial inner membrane potential, a key readout for mitochondrial general fitness. Using ΔΨm-sensitive tetramethylrhodamine methyl ester (TMRM) probe, we showed that *SYBU* deficiency is associated with a significant decrease in mitochondrial membrane potential, which is further reduced upon PTX treatment (Fig. 3G, Suppl Fig. S4J), consistent with reduced mitochondrial depolarization and impaired mitochondrial respiration. Reduced mitochondrial dependence in *SYBU*-depleted cells was also observed using the SCENITH™ technology (Single-Cell Metabolism by Profiling Translation Inhibition) that analyzes the metabolic profiles of cell populations by flow cytometry [31] (suppl Fig. S4K). Together these results indicate that *SYBU* depletion and PTX treatment alter mitochondrial respiratory activity with no detectable effect on mitochondrial abundance, consistent with a role in function rather than biogenesis. These observations are consistent with recent work showing that a low dose of Taxol impairs oxidative phosphorylation, depolarizes mitochondria and promotes a shift toward glycolysis in actively respiring cancer cells [32], supporting the view that taxanes can act as mitochondrial stressors in addition to their effects on spindle microtubules.

Measurement of extracellular acidification rate (ECAR) revealed that glycolysis is significantly increased upon *SYBU*-depletion and further enhanced by PTX treatment of COV318 cells (Fig. 3H, Suppl Fig. S4L). In line with seahorse experiments, SCENITH flow cytometry analysis of cell metabolism showed increased glycolytic capacity in *SYBU*-depleted cells compared to control (suppl Fig. S4M). Together, these results indicate that *SYBU* depletion and PTX treatment favor a metabolic switch from mitochondrial respiration to glycolysis, consistent with a Warburg effect. The metabolic switch induced by PTX treatment is more pronounced upon *SYBU*-depletion, further indicating that *SYBU*-deficient cells are more sensitive than control ones to metabolic plasticity induced by chemotherapy.

### SYBU depletion alters mitochondrial morphology and cristae ultrastructure

Mitochondrial function and structure being closely interlinked [33–35], we examined mitochondrial ultrastructure by transmission electron microscopy (TEM) upon *SYBU* depletion (Fig. 4A). Mitochondrial morphology was affected in *SYBU*-depleted COV318 cells with a significant reduction in the percentage of elongated mitochondria (Fig. 4B). The mean surface and perimeter of mitochondria were reduced, whereas their circularity was increased, in *SYBU*-depleted cells compared to control (suppl Fig. S5A-S5C). Mitochondrial aspect ratio, that reflects the networking aspect of mitochondria, was also decreased in *SYBU*-depleted cells (Suppl Fig.S5D). Under our experimental conditions, treatment with PTX had no significant effect on mitochondrial morphology, both in control and *SYBU*-deficient cells (Fig. 4B). In contrast, both *SYBU* depletion and PTX treatment induced marked alterations in mitochondrial cristae architecture (Fig. 4C, 4D). In control cells, mitochondria displayed a typical ultrastructure characterized by uniformly distributed, lamellar cristae extending perpendicularly from the inner mitochondrial membrane (IMM). By contrast, both *SYBU*-depleted and PTX-treated COV318 cells exhibited severe disorganization of the IMM, accompanied by irregular cristae structures and mitochondrial regions partially devoid of cristae. *SYBU*-depletion exacerbated the deleterious effect of PTX on cristae architecture (Fig. 4D) which is crucial for mitochondrial function, indicating that *SYBU*-depleted cells are more sensitive to chemotherapeutic treatment on cristae organization.

**Figure 4.**
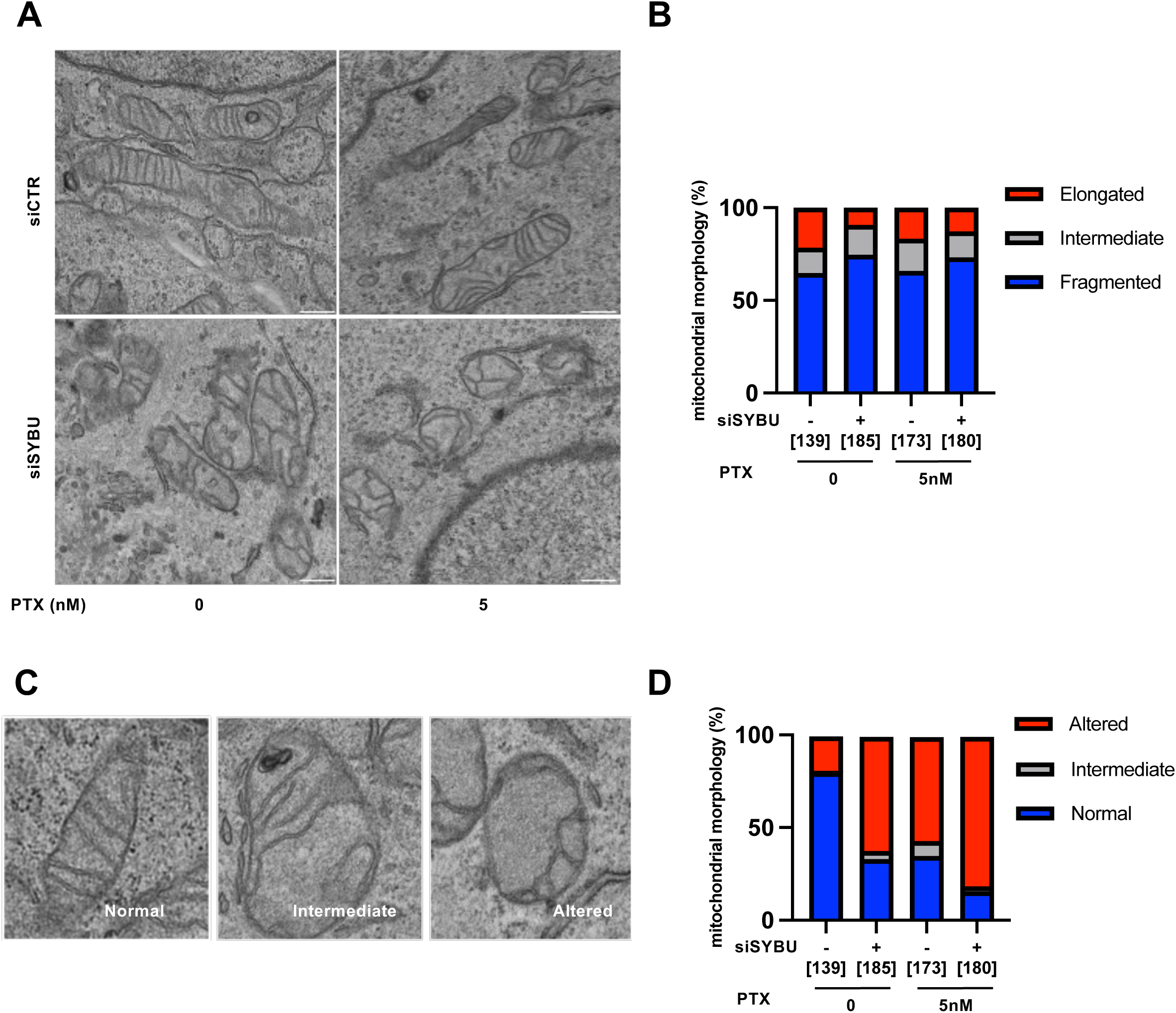
*SYBU* is required for structural organization of mitochondria cristae. A. Representative transmission electron microscope (TEM) images of COV318 cells expressing (siCTR) or not (siSYBU) *SYBU*, treated or not with 5nM PTX during 72 hrs. Scale bar 10μm. B. Percentage of mitochondrial morphology types on images in (A). C. Representative TEM images of COV318 cells with different cristae phenotypes. D. Percentage of different cristae phenotypes as in (C) from images of COV318 cells treated as in (A). (B, D). The number of mitochondria analyzed is in brackets.

### Syntabulin associates with VDAC1 and MIC60 at mitochondria

To get insight into the molecular mechanisms regulated by syntabulin in ovarian cancer cells, a proteomic approach was undertaken to identify its interactome. COV318 cell lysates expressing either GFP or GFP-SYBU fusion protein were immunoprecipitated using anti-GFP antibodies and molecular complexes were analyzed by mass spectrometry. A total of 301 quantified proteins that were selectively retained in GFP-SYBU precipitates were identified, 52 of which were selected for further analysis as they reached high confidence (Suppl Table TS6). Syntabulin partners belong to four major functional networks centered on mitochondrial membrane organization, microtubule cytoskeleton, ubiquitination and protein folding (Fig. 5A, Suppl Fig. S6A, S6B).

**Figure 5.**
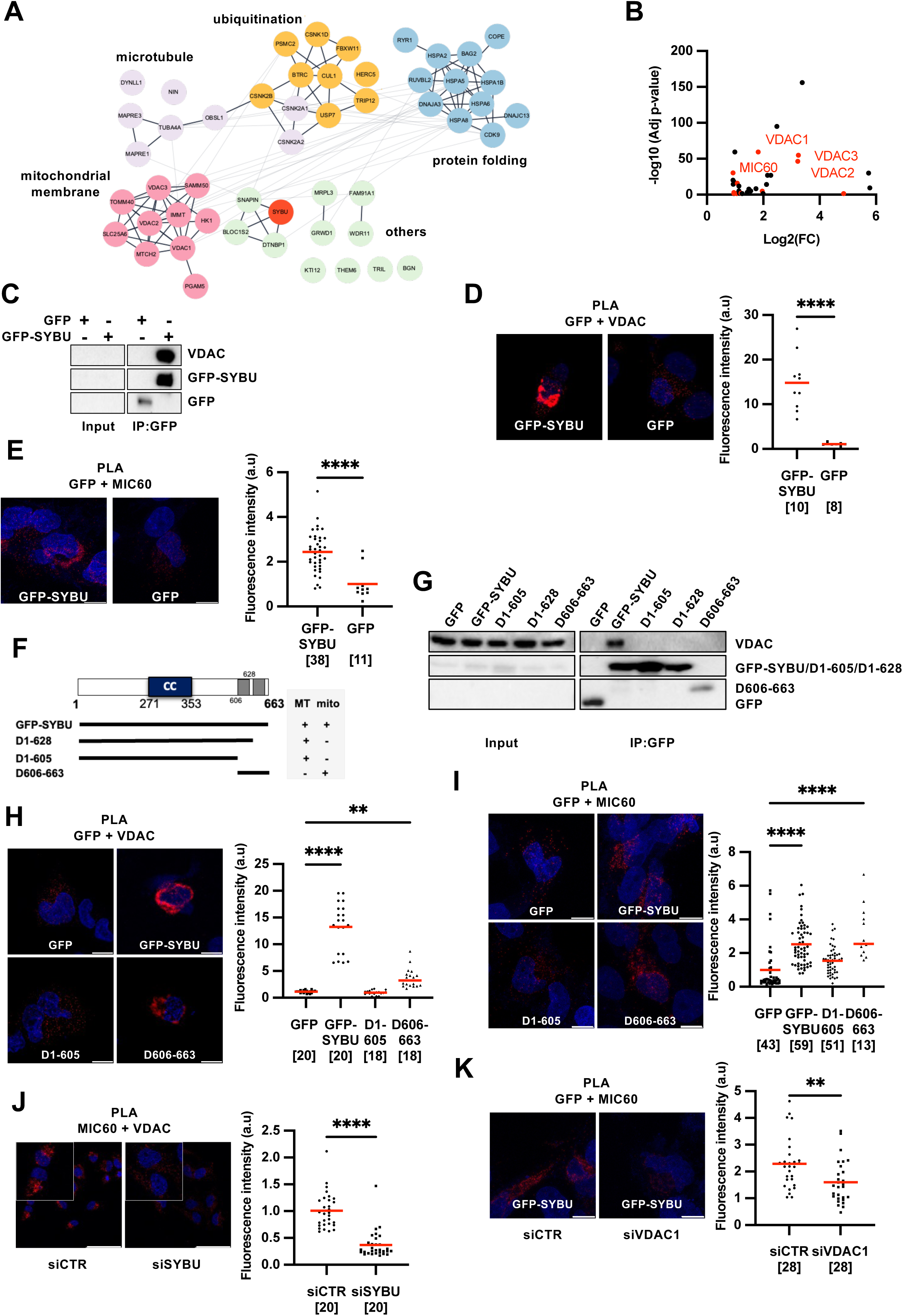
Syntabulin interacts with VDAC1 and MIC60. A. Protein-protein interaction (PPI) network of SYBU-interacting proteins reaching high confidence in the quantitative label-free mass spectrometry analysis of COV318 cells. Enrichment analysis was visualized by using Cytoscape software. B. Volcano plot showing distribution of SYBU-interacting proteins reaching high confidence in the quantitative label-free mass spectrometry analysis of COV318 cells. Mitochondrial proteins with highest p value are indicated in red. x axis = log2(fold-change), y axis = − log10(p-value). C. Immunoprecipitation assay of COV318 cell lysates expressing GFP-SYBU or GFP. Blots were probed with anti-VDAC antibodies and reprobed with anti-GFP antibodies to reveal GFP-SYBU and GFP. D. COV318 cells were transfected with GFP or GFP-SYBU constructs and analyzed by proximity ligation assay (PLA) using mouse anti-GFP and rabbit anti-VDAC1 primary antibodies. Representative images (left) and quantification of PLA intensity signals (a.u, arbitrary units) per region of interest (right panel). E. COV318 cells were transfected as in (D) and analyzed by PLA using rabbit anti-GFP and mouse anti-MIC60 primary antibodies. Representative images (left) and quantification of PLA intensity signals as in (D) (right panel). F. Schematic representation of the SYBU protein sequence illustrating the position of CC: coiled coil region and transmembrane regions (in grey). SYBU deletion mutants and their ability (+) or not (−) to bind microtubules (MT) or mitochondria (mito) are shown below. Amino acid numbering is from accession number NP_001093224.1. G. Immunoprecipitation assay of COV318 cell lysates expressing GFP, GFP-SYBU or GFP-fused D1-605, D1-628 or D6606-663 deletion mutants. Blots were probed with anti-VDAC and anti-GFP antibodies to reveal VDAC and GFP-fused SYBU and deletion mutants as indicated. H. COV318 cells were transfected with GFP or GFP-fused constructs as indicated and analyzed by PLA using mouse anti-GFP and rabbit anti-VDAC primary antibodies. Representative images (left) and quantification (right) as in (D). I. COV318 cells were transfected as in (H) and analyzed by PLA using rabbit anti-GFP and mouse anti-MIC60 primary antibodies. Representative images (left) and quantification (right) as in (D). J. COV318 cells were transfected with control (siCTR) or SYBU-targeting siRNA (siSYBU) and analyzed by PLA using rabbit anti-VDAC1 and mouse anti-MIC60 primary antibodies. Representative images (left) and quantification (right) as in (D). K. COV318 cells were transfected with GFP-SYBU and control (siCTR) or VDAC1-targeting siRNA (siVDAC1) and analyzed by PLA using rabbit anti-GFP and mouse anti-MIC60 primary antibodies. Representative images (left) and quantification (right) as in (D). (D, E, H-K) Number of cells is indicated in brackets. \*\**P* < 0.01, \*\*\*\**P* < 0.0001.

Given the functional impact of syntabulin on mitochondrial metabolism, we focused our attention on the mitochondrial interactome. Porins (Voltage-dependent anion-selective channel proteins) VDAC1, VDAC2 and VDAC3 were among mitochondrial partners with highest significance in the mass spectrometry analysis (Fig. 5B, suppl Table TS6). Interaction between syntabulin and VDAC was validated in co-immunoprecipitation experiments (Fig. 5C). Molecular proximity between syntabulin and VDAC1 in ovarian cancer cells *in situ* was further confirmed by proximity ligation assay (PLA) (Fig. 5D, suppl Fig. S6C). Interestingly MIC60, a protein of the mitochondrial inner membrane playing a central role in the organization of cristae [36–38] was also identified among syntabulin-interacting proteins with highest significance in the mass spectrometry analysis (Fig. 5B, Suppl Table TS6). Molecular proximity between syntabulin and MIC60 was confirmed *in situ* in PLA experiments (Fig. 5E).

Syntabulin is a 663 amino acids protein that presents a carboxy-terminal region with two alpha helices spanning the mitochondrial outer membrane, a central coiled-coil domain and an intrinsically disordered cytosolic amino-terminal region that binds microtubules (Fig. 5F). When expressed in ovarian cancer cells, syntabulin co-localizes with mitochondria and decorates the microtubule cytoskeleton (Suppl Fig. S6D). Deletion mutants were constructed to identify interacting domains of the protein. Constructs D1-628 and D1-605 are devoid of the C-terminal alpha helices anchored in the mitochondrial outer membrane (Fig. 5F). As expected, these fusion proteins decorate microtubules but do not colocalize with mitochondria (Suppl Fig. S6D). Deletion mutant D606-663, that contains the C-terminal alpha helices but lacks the N-terminal part, co-localizes with mitochondria but not with the microtubule cytoskeleton (Fig. 5F, Suppl Fig. S6D). Co-immunoprecipitation and PLA experiments indicate that the N-terminal region D1-605 of syntabulin is not sufficient to maintain its association with VDAC1 and MIC60, whereas the C-terminal mitochondrial transmembrane region (D606-D663) retains close proximity to both proteins (Fig. 5G-5I). Thus, the C-terminal mitochondrial targeting domain of syntabulin is required for its association with both VDAC1 and MIC60.

Previous studies have revealed the existence of a molecular complex between VDAC1 and MIC60, although these proteins are localized in the outer and inner mitochondrial membrane, respectively [39]. We confirmed molecular proximity between endogenous VDAC1 and MIC60 in COV318 cells (Fig. 5J; suppl Fig. S7A, S7B). We then investigated whether syntabulin may be part of this complex by silencing *SYBU* prior to analyzing VDAC1/MIC60 complexes by PLA. A marked reduction in the intensity of PLA signals generated by endogenous VDAC1 and MIC60 proteins (Fig. 5J) indicates that syntabulin indeed contributes to this interaction. On the other hand, depleting VDAC1 in COV318 cells also led to a significant reduction of the PLA signals formed between syntabulin and MIC60 (Fig. 5K), indicating that VDAC1 contributes to the association between syntabulin and MIC60. Together, these findings support a model in which syntabulin, via its C-terminal mitochondrial anchor, associates with VDAC1 at the outer membrane and contributes to VDAC1–MIC60 coupling across the outer and inner mitochondrial membranes, thereby providing a structural link that can influence cristae organization and mitochondrial function.

### A Syntabulin-MIC60 axis couples mitochondria ultrastructure with mitotic fidelity

We next asked whether syntabulin, by stabilizing the interaction between VDAC1 and MIC60, may influence the abundance of individual components of the complex. Western blot analysis of *SYBU*-deficient COV318 cancer cells revealed that loss of *SYBU* induces a marked decrease in MIC60 protein levels with no major change in VDAC protein levels (Fig. 6A). Given that MIC60 is a key determinant of cristae architecture, and that *SYBU* depletion leads to profound cristae disorganization, we reasoned that impaired cristae structures observed upon *SYBU* silencing in ovarian cancer cells may be a consequence of decreased MIC60 protein levels. To evaluate this hypothesis, a GFP-MIC60 fusion protein was expressed into *SYBU*-deficient cells, in comparison to GFP and GFP-SYBU as controls, and mitochondrial ultrastructure was analyzed by electron microscopy. Results showed that expressing GFP-MIC60 in *SYBU*-depleted COV318 cells restored mitochondria elongation and cristae organization at a level similar to that achieved with GFP-SYBU expression (Fig. 6B-6D; Suppl Fig. S7C-S7F), suggesting that the effects of syntabulin on cristae structure indeed involve MIC60. Furthermore, the rescue of mitochondrial membrane potential TMRM following GFP-MIC60 expression in *SYBU*-depleted COV318 cells, to the same level as GFP-SYBU (Fig. 6E), supports a central role for the syntabulin–MIC60 axis in sustaining mitochondrial function.

**Figure 6.**
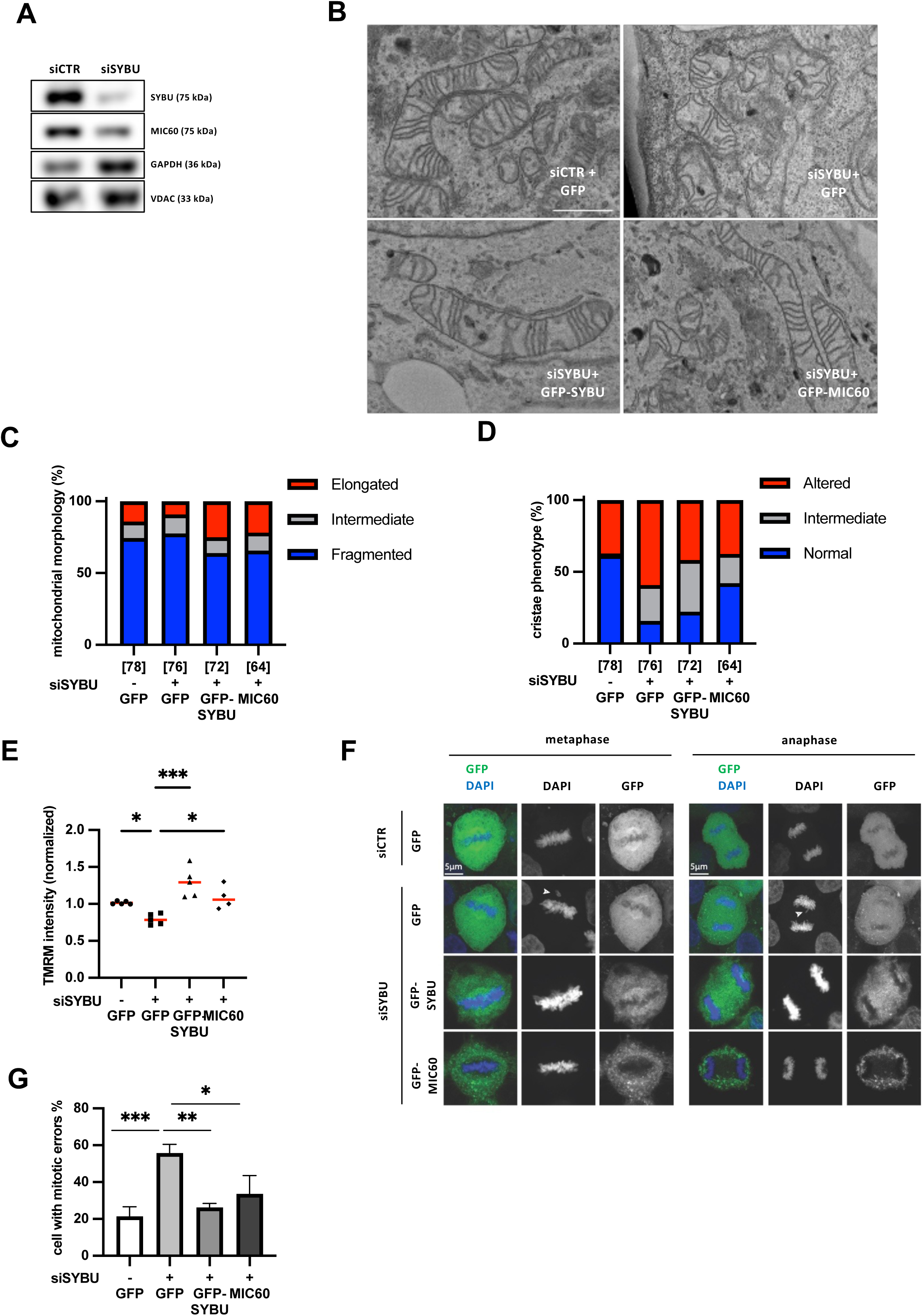
A Syntabulin-MIC60 axis couples mitochondria ultrastructure with mitotic fidelity. A. Representative immunoblot showing abundance of 3 mitochondrial proteins (SYBU, MIC60, and VDAC) in COV318 cells expressing (siCTR) or not (siSYBU) endogenous SYBU. GAPDH is used as internal control of the amount of total proteins. Molecular weight is indicated in kilodalton (kDa) on the right. B. Representative transmission electron microscope (TEM) images of COV318 cells expressing (siCTR) or not (siSYBU) endogenous SYBU and transfected with GFP, GFP-SYBU or GFP-MIC60 as indicated. Scale bar 10μm. C. Percentage of mitochondrial morphology types based on TEM images in (B). D. Percentage of different cristae phenotypes based on TEM images in (B). E. Quantification of relative TMRM fluorescence intensity of COV318 cells expressing (-) or not (siSYBU) endogenous SYBU and transfected with GFP, GFP-SYBU or GFP-MIC60 as indicated. F. Immunofluorescence photographs of HeLa cells expressing (siCTR) or not (siSYBU) endogenous SYBU and transfected with GFP, GFP-SYBU or GFP-MIC60 prior staining with anti-GFP and DAPI. Magnification, 63x. Scale bar 5μm. G. Percentage of HeLa cells with mitotic errors shown in (F). \**P* < 0.05; \*\**P* < 0.01; \*\*\**P* < 0.001.

Strikingly, expression of GFP-MIC60 into SYBU-deficient cells also proved sufficient to restore the mitotic phenotype characterized by chromosome misalignments in metaphase and chromosome missegregation in anaphase, to the same extent as GFP-SYBU (Fig. 6F, 6G). These results highlight the link between cristae organization, mitochondrial metabolism and mitotic integrity.

Taken together, our data suggest that intact mitochondrial cristae are important to ensure correct chromosome alignment and segregation during mitosis and reveal the crucial contribution of the syntabulin–MIC60 axis to coupling mitochondrial ultrastructure with mitotic fidelity.

## DISCUSSION

The search for molecular markers of HGSOC chemoresistance is an unmet medical need. In this study, we identify *SYBU* as a potential predictive biomarker of chemoresistance to taxane-based chemotherapy. *SYBU* is down-regulated in cancer tissues and its low levels are associated with agressivity of the tumor. *SYBU* encodes syntabulin, a 663 amino-acid polypeptide of the mitochondrial outer membrane that binds microtubules. *SYBU* depletion in HGSOC cell lines causes major defects in mitochondria ultrastructure, with disorganization of the cristae, associated with marked reduction in mitochondrial respiration compensated by elevated glycolysis, a metabolic switch known as the Warburg effect. *SYBU* depletion also leads to severe mitotic defects including centrosome amplification, chromosome misalignment in metaphase and missegregation in anaphase, which are recognized sources of aneuploidy.

Treatment of *SYBU*-depleted cells with paclitaxel increases cristae alterations and metabolic rewiring, and further exacerbates mitotic defects, ultimately leading to cancer cell death. Both metabolic reprogramming and aneuploidy are hallmarks of cancer that can promote tumor evolution and aggressive behaviour [40,41]. However, several studies indicate that when chromosome missegregation rates exceed a tolerable threshold, extreme chromosomal instability (CIN) becomes detrimental and sensitizes tumors to microtubule-targeting agents such as taxanes [14,22–24]. This is consistent with our data on ovarian cancer patients showing that *SYBU*-deficient tumors are of poor prognosis, yet aggressive *SYBU*-deficient ovarian tumors are also more sensitive to taxane-based chemotherapy.

We propose a model in which in *SYBU*-deficient cells, taxanes markedly reduce mitochondrial respiration and increase mitotic defects, so that the combination of mitochondrial dysfunction and prominent mitotic errors accumulates cellular abnormalities beyond a tolerable threshold. *SYBU*-deficient cells being defective in two important cellular functions, namely mitochondrial metabolism and cell division, are therefore more vulnerable to chemotherapy (Fig. 7).

**Figure 7.**
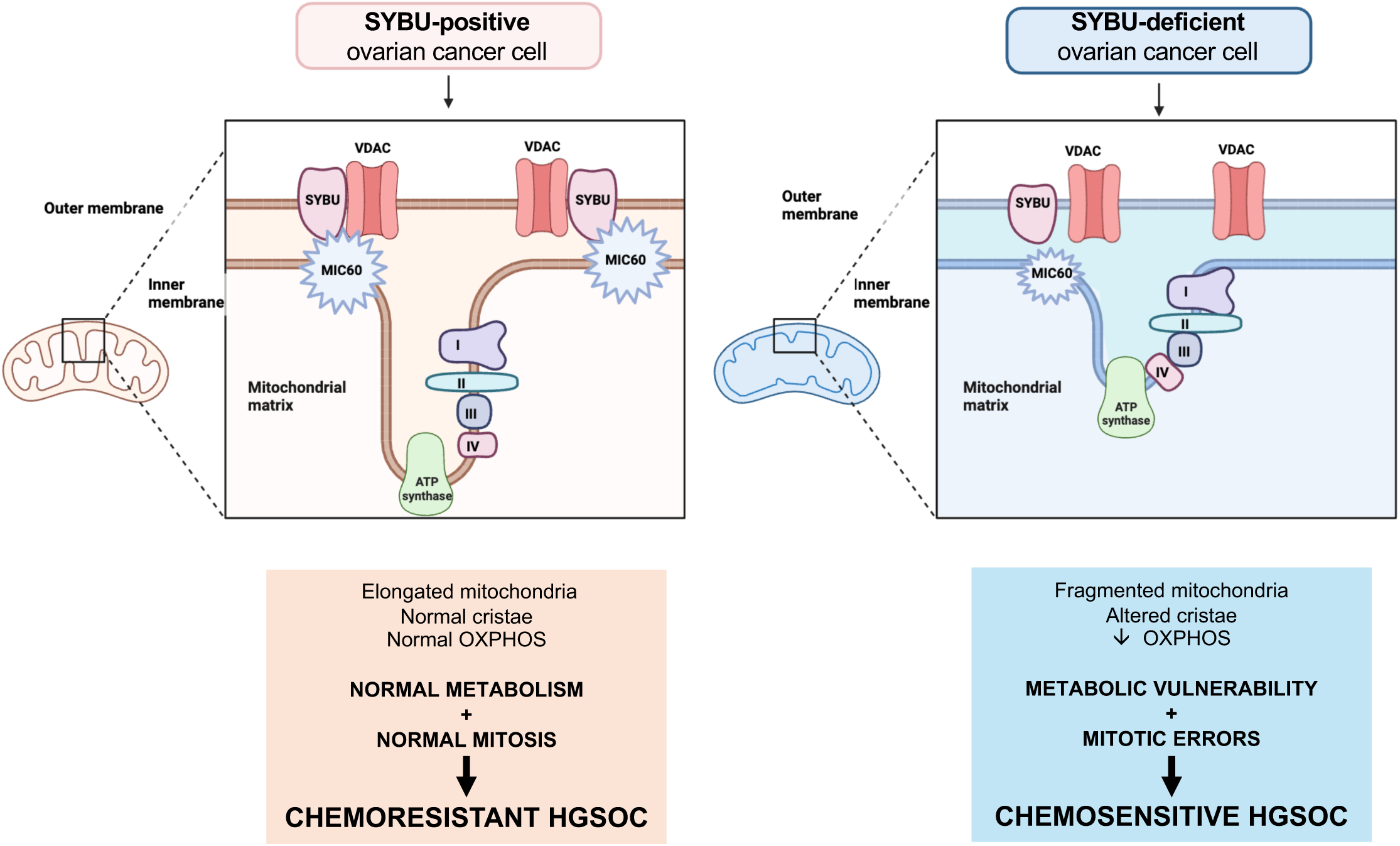
Graphical abstract. In *SYBU*-positive HGSOC cancer cells (left), SYBU interacts with VDAC and MIC60 to maintain cristae integrity, normal metabolism and mitotic fidelity. In SYBU-low cancer cells (right), MIC60 levels are decreased, the VDAC-MIC60 complex is disrupted and mitochondrial cristae are altered. This causes defects in mitochondrial metabolism and mitotic features, which are two major sources of vulnerability to chemotherapy.

Syntabulin is a mitochondrial outer membrane protein that interacts with the VDAC1 porin, a well-known regulator of mitochondrial function. Syntabulin also contributes to VDAC1 association with the mitochondrial inner membrane protein MIC60, a major component of the MICOS complex that controls the organization of cristae. We show here that syntabulin controls VDAC-MIC60 interaction. Its depletion in HGSOC cells triggers a marked reduction in protein levels of MIC60, with subsequent alterations in mitochondrial cristae, which are the principal sites of energy production by oxidative phosphorylation. Overexpression of MIC60 in *SYBU*-deficient cancer cells restores cristae integrity, mitochondrial functions as well as mitotic fidelity, pointing to the existence of a syntabulin-MIC60 axis controlling HGSOC homeostasis.

The microtubular localization of syntabulin, together with its impact on the time spent in mitosis, centrosome number and chromosome alignment, suggests that *SYBU* contributes to a mitochondria–microtubule interface that coordinates mitochondrial architecture with mitotic features. A bidirectional relationship between mitochondrial functions and mitotic progression has been highlighted, although the mechanistic links are not fully understood [42]. Mitochondria are highly dynamic organelles that undergo coordinated waves of fission and fusion across the cell cycle, and several studies have linked these dynamics and organelle redistribution to mitotic progression and spindle function [43–45]. Our findings place syntabulin at the interface between mitochondria and the microtubule cytoskeleton. In neurons, syntabulin has been shown to mediate anterograde transport of mitochondria along microtubules, ensuring proper organelle positioning in axons and synapses [18,19]. Recent studies from our group have revealed that syntabulin also controls mitochondria transport and positioning in migrating breast cancer cells [20]. Organelle positioning and fission patterns are increasingly recognized as critical for mitotic spindle function, chromosome segregation and organelle inheritance. Recent work has also disclosed general principles of organelle positioning, showing that unequal segregation of mitochondria during asymmetric divisions can drive fate divergence and impact genome stability *in vivo* [46,47]. Whether the disruption of cristae organization caused by *SYBU* down-regulation in cancer cells leads to reduced mitochondrial bioenergetics around mitotic structures, thereby weakening spindle function and/or increasing chromosome segregation errors, warrants further investigation.

From a clinical perspective, our data identify the syntabulin–MIC60 axis as a nexus between mitochondrial metabolism, cristae architecture and mitotic fidelity that shapes the response to taxane-based chemotherapy. Metabolic heterogeneity has emerged as a key determinant of response to chemotherapy in ovarian cancer, with OXPHOS-high tumors displaying enhanced sensitivity to platinum–taxane regimens and vulnerability to OXPHOS inhibition [48,49]. Targeting OXPHOS in HGSOC to eliminate stem-like cancer cells has recently been reported [50]. In this metabolic landscape, low *SYBU* expression defines a state characterized by impaired OXPHOS, increased glycolytic reliance and heightened sensitivity to taxane-induced mitochondrial stress, suggesting that *SYBU* status and associated mitochondrial phenotypes could be incorporated into future stratification strategies.

From a therapeutic point of view, targeting the syntabulin-MIC60 axis in high-*SYBU* chemoresistant HGSOC may potentially sensitize tumors to chemotherapy by disrupting the mitochondrial molecular complex. In the light of precision medicine, these findings may have important implications for HGSOC patients, pointing to *SYBU*-associated pathways as potential targets to improve therapeutic outcomes by overcoming chemoresistance, especially in combination with agents that impact microtubules or mitochondrial function.

## METHODS

### Patients and samples

The CURIE cohort of patients (GSE 26193) has been previously described [48,51,52]. All patients signed an informed consent for voluntary participation in the trial. Studies were approved by the scientific committee of the Institut Curie. Only patients (n=42) with high grade serous ovarian cancer who received taxane-based chemotherapy were included in our study. RNA extraction and profiling on Affymetrix U133 Plus 2.0 GeneChips, as well as microarray data normalization were performed as described [51].

The AOCS (Australian Ovarian Cancer Study) cohort of patients (GSE 9899) was described previously [53]. Only patients (n=136) with high grade serous cancer treated with taxane-based chemotherapy were included in our study. Clinical data for patients of both cohorts are presented in Supplementary Table S2. Patients were considered as sensitive (S) if they have not relapsed within 12 months after the end of the chemotherapy cycle. Patients considered as resistant (R) to chemotherapy were those who relapsed less than twelve months after diagnosis. Progression-free survival curves of (S) and (R) patients in each cohort are presented in Suppl Fig.S1A.

The GR cohort (E-MTAB-11177) comprises a total of 39 patients with ovarian cancer, including 20 borderline tumors (BOL) and 19 adenocarcinomas (ADK). Patients were recruited at the Gustave Roussy Hospital (Villejuif, France). Samples were analyzed by pathologists. Patients were informed of the research study and signed a consent for voluntary participation. The study was approved by the scientific committee of the Institut Gustave Roussy.

### Transcriptomic analysis

For CURIE and AOCS cohorts, analysis was carried out on expressed data from the GSE26193 matrix.txt.gz (https://www.ncbi.nlm.nih.govgeo/query/acc.cgi?acc=GSE26193) and GSE9899 series matrix.txt.gz (https://www.ncbi.nlm.nih.gov/geo/query/acc.cgi?acc=GSE9899). Affymetrix U133 Plus 2 data were imported in BrB Array Tools Version: 4.5.1 (August 2016) [https://brb.nci.nih.gov/BRB-ArrayTools]. Normalized data were used for a Class Comparison between resistant and sensitive samples. For the GR cohort, following RNA extraction, gene expression studies were performed in dual color on Agilent Whole Human Genome Oligo Microarray A-AGIL11 (design 012391). Detailed sample information is available at https://www.ebi.ac.uk/biostudies/arrayexpress/studies/E-MTAB-11177. Genes that were differentially expressed among the two compared classes of samples were identified using a random-variance t-test. Genes were considered statistically significant if their P value was <0.05. A stringent significance threshold was used to limit the number of false-positive findings and an FDR adjustment (0.05) was used for multiple testing.

### Cell Lines

The human epithelial HGSOC cell line COV318 was kindly provided by Dr Fatima Mechta-Grigoriou (Institut Curie, Paris, France). HeLa cells were provided by Dr. Mounira Amor-Gueret (Institut Curie, Orsay, France). HeLa-H2B cells stably expressing mCherry-Histone H2B were kindly provided by Dr Valérie Doye (Institut Jacques Monod, Paris, France). All cell lines were grown in Dulbecco’s modified Eagle’s medium (DMEM, GIBCO, Thermo Fisher Scientific) containing 10% fetal bovine serum (FBS, GIBCO, Thermo Fisher Scientific). Cells were cultured at 37°C and 5% CO2 and were split every 3-4 days. They were routinely authenticated by morphologic observation and tested for absence of mycoplasma contamination using VenorGeM Advance detection kit (Minerva Biolabs).

Paclitaxel (Fresenius Kabi, France) was added to the cell media from concentrated stock (7mM) dissolved in sterile media culture.

### Plasmid constructs

GFP-SYBU (also designated FL) plasmid was obtained by subcloning the full-length SYBU cDNA insert (accession number NP_001093214) of SYBU-GFP (Origene RG217887) into the HindIII-KpnI cloning sites of pEGFP-C1 vector (Clontech, CA, USA). Deletion mutants D1-628 and D1-605 were obtained from the *SYBU* sequence by using QuikChange® II XL Site-Directed Mutagenesis Kit (Agilent) with following oligonucleotides:

(5’-CGGTTCTGTGGGCATTCtaaGGTACCGCGGGCCCGG-3)’and

(5’ GGCAGTACTGGAGCAGCtaaGGTACCGCGGGCCC-3’), respectively.

SYBU mutant D606-663 was obtained by subcloning the PCR product of oligonucleotides: sens (5’-attAAGCTTtcAGCTTCCTGGTGGATCTCC-3’), and anti-sens (5’-aatGGTACCttaGGTTTTGATACGGAAGGCGGTG −3’), into the HindIII-KpnI cloning sites of pEGFP-C1 vector.

For rescue experiments, all constructs were made resistant to siRNA by using QuikChange® II XL Site-Directed Mutagenesis Kit and oligonucleotide sequence: (5’-CTGGAAGGTACATGTCCTGTGGAGAAAATCATGGTGGTC-3’).

### Transfections

Small interfering RNA (siRNA) was transfected at 20nM for 48 hrs or 72 hrs by using Lipofectamine Reagent 2000 (Invitrogen) as described by the manufacturer. Control siRNA duplex (non-targeting pool #1), and specific *SYBU* siRNA (on-target plus siRNA sens strand GUA CAU GUC UUG CGG UGAA), *SYBU* siRNA#2 (on-target plus siRNA sens strand GAG UAA UGG AGC UUC GUCA), VDAC1 siRNA (on-target plus siRNA sens strand UAA CAC GCG CUU CGG AAUA) were purchased from Dharmacon (Chicago, IL, USA). Silencing efficiency was measured by Western Blot and real time RT-PCR. For *SYBU* knockdown the following shRNA (GCGTGGAAGAGAGGTTGGA) was purchased from GE Healthcare (France).

### Mitotic error quantification

Cells were synchronized at various stages of mitosis using the double Thymidine block and release protocol (DTBR). Initially, cells were treated with 2 mM Thymidine twice for 16hrs, followed by three washes with warm media and an 8hrs incubation period without Thymidine. Subsequently, following the completion of the second Thymidine block, cells were released into Thymidine-free media and collected 10hrs after release, corresponding to the mitotic peak. In specific experiments, Paclitaxel was added 5h after release from Thymidine treatment at concentrations of 2 nM, followed by a 5hrs incubation period prior to cell collection. At least 100 cells were counted per condition and per experiment. The presence of mitotic errors was assessed by DAPI staining based on the morphology of the cell and cells were categorized in those that had normal mitotic shape and those that displayed different types of mitotic defects (chromosome misalignments, lagging chromosomes, and/or anaphase/telophase bridges). The number of centrosomes was assessed by staining with the centrosome marker Pericentrin and cells were categorized in those with normal (2) or multiple (>2) centrosomes. Images were taken with a 63X objective using confocal microscopy (Leica).

### Live cell imaging

For live cell imaging, HeLa-mCherryH2B cells were transfected for 48 hrs with control-siRNA or specific *SYBU* siRNA and incubated with siR-Tubulin dye (2.5 nM) prior treatment with PTX (2 nM). Cells were imaged on a confocal laser scanning microscope TCS SP8 MP (Leica), with 40X objective, every 8 min for 60 hrs. Multi-dimensional acquisitions were performed in time-lapse mode using ICy software. Image analysis was performed using ImageJ software.

### Co-immunoprecipitation

COV318 cells were transfected for 24 hrs with appropriate GFP-tagged plasmid using X-tremeGENE 9 (Roche) and lysed in RIPA lysis buffer (0.2 M PMSF, 1mM aprotinin, 5mg/ml leupeptin, 2mg/ml pepstatin, 0.2M sodium orthovanadate, 1M sodium fluoride, and 10μM okadaic acid). Lysates were centrifuged, incubated with uncoupled magnetic agarose beads at 4°C for 1min, and then incubated with anti-GFP VHH coupled to magnetic agarose beads (GFP-trap_MA, Chromotek) for 2 hrs at 4°C. Antibody beads were recovered with a bar magnet, washed three times with PBS, and analyzed by western blotting with appropriate antibodies. Results shown are representative of 3-5 independent experiments.

### Proximity ligation assay (PLA)

PLA detection *in situ* was performed with Duolink II In Situ Far Red kit (Sigma-Aldrich, St Louis, USA). Briefly, COV318 cells were transfected for 24 hrs with GFP-tagged plasmids or 48h with siRNA, then fixed with paraformaldehyde 4% and permeabilized with Triton X100 0,1%. Primary antibodies anti-MIC60 (Abcam, ab110329), anti-VDAC (Abcam, ab15895) and anti-GFP (Abcam, ab1218) were incubated over night at 4°C. Slides were then incubated for 1 hr at 37°C with PLA DUOLINK probes anti-mouse PLUS and anti-rabbit MINUS (1:5) and treated for ligation and amplification with oligonucleotide probe associated to Cyanine 5 red fluorescent dye according to the supplier.

Images were acquired on Leica SP8 STED confocal microscope with a 63X immersion lens with the Leica Application Suite X software. PLA intensity signals per region of interest (ROI) were quantified with the « polyline » tool of this software in the « quantify » menu.

### Proteomics and mass spectrometry

CO318 cells were transiently transfected with 2 µg of SYBU-GFP or GFP plasmids for 24 hrs. For immuno-isolated samples, 1 mg extract was incubated with magnetic beads (GFP-trap_MA; Chromotek) for 2 hrs at 4 °C. Proteins on magnetic beads were washed twice with 25 mM NH_4_HCO_3_ and processed as described before [54]. On-beads digestions were performed by adding 0.2 μg of trypsin-LysC (Promega) for 1 hr at 37°C. Samples were then loaded into custom-made C18 StageTips packed by stacking one AttractSPE® disk (#SPE-Disks-Bio-C18-100.47.20 Affinisep) and 2mg beads (#186004521 SepPak C18 Cartridge Waters) into a 200 µL micropipette tip for desalting. Peptides were eluted using a ratio of 40:60 CH_3_CN:H_2_O + 0.1% formic acid and vacuum concentrated to dryness with a SpeedVac apparatus. Peptides were reconstituted in injection buffer in 0.3% trifluoroacetic acid (TFA) before liquid chromatography-tandem mass spectrometry (LC-MS/MS) analysis.

#### LC-MS/MS Analysis

Online chromatography was performed using an RSLCnano system (Ultimate 3000, Thermo Fisher Scientific) coupled to an Orbitrap Exploris 480 mass spectrometer (Thermo Fisher Scientific). Peptides were trapped on a C18 column (75 μm inner diameter × 2 cm; nanoViper Acclaim PepMap 100, Thermo Fisher Scientific) with buffer A (2:98 CH_3_CN:H2O in 0.1% formic acid) at a flow rate of 3.0 µl/min over 4 min. Separation was performed using a 50 cm × 75 μm C18 column (nanoViper Acclaim PepMap RSLC, 2 μm, 100 Å), regulated to a temperature of 50 °C with a linear gradient of 2% to 25% buffer B (100% CH_3_CN in 0.1% formic acid) at a flow rate of 300 nl min^−1^ over 91 min. MS full scans were performed in the ultrahigh-field Orbitrap mass analyzer in the m/z range of 375–1500 with a resolution of 120,000 at m/z 200, a automatic gain control (AGC) set at 300% and with a maximum injection time (IT) of 25 ms. The 20 most intense ions were isolated (isolation width of 1.6 m/z) and further fragmented via high-energy collision dissociation (HCD) activation and a resolution of 15,000, an AGC target value set to 100% and with a maximum IT of 60s. We selected ions with charge state from 2+ to 6+ for screening. Normalized collision energy (NCE) was set at 30 and a dynamic exclusion of 40 seconds.

#### Data analysis

For identification, the data were searched against the Homo sapiens UP000005640 database (downloaded 12/2019 containing 20364 entries) using Sequest HT through Proteome Discoverer (version 2.4). Enzyme specificity was set to trypsin and a maximum of two missed cleavage sites were allowed. Oxidized methionine, N-terminal acetylation, methionine loss and methionine acetylation loss were set as variable modifications. Maximum allowed mass deviation was set to 10 ppm for monoisotopic precursor ions and 0.02 Da for MS/MS peaks. The resulting files were further processed using myProMS [55] [https://github.com/bioinfo-pf-curie/myproms] v.3.9.3. False-discovery rate (FDR) was calculated using Percolator [56] and was set to 1% at the peptide level for the whole study. Label-free quantification was performed using peptide extracted ion chromatograms (XICs), computed with MassChroQ v.2.2.1[57]. For protein quantification, XICs from proteotypic peptides shared between compared conditions (TopN matching) with missed cleavages were used. Median and scale normalization at peptide level was applied on the total signal to correct the XICs for each biological replicate (N=5). To estimate the significance of the change in protein abundance, a linear model (adjusted on peptides and biological replicates) was performed, and p-values were adjusted using the Benjamini–Hochberg FDR procedure. Proteins with at least three total peptides in all replicates (n=5), a 1.9-fold enrichment and an adjusted p-value ≤ 0.05 were considered significantly enriched in sample comparisons. Unique proteins were considered with at least two total peptides in three replicates. A total of 52 proteins selected with these criteria were further analyzed or subjected to GO functional enrichment analysis.

### Transmission electron microscopy

Samples were fixed with 2% glutaraldehyde in 0.1 M Na cacodylate buffer pH 7.2 and then contrasted with Oolong Tea Extract (OTE) 0.2% in cacodylate buffer, postfixed with 1% osmium tetroxide containing 1.5% potassium cyanoferrate, gradually dehydrated in ethanol (30% to 100%) and substituted gradually in mix of ethanol-Epon and embedded in Epon from Delta microscopies (France). Thin sections (70 nm) were collected onto 200 mesh copper grids, and counterstained with lead citrate. Grids were examined with Hitachi HT7700 electron microscope operated at 80kV from Milexia (France), and images were acquired with a charge-coupled device camera (AMT) in MIMA2 Microscopy and Imaging Facility for Microbes, Animals and Foods https://doi.org/10.15454/1.5572348210007727E12.

Mitochondrial and cristae phenotype quantification was obtained using Image J by manually tracing mitochondria from TEM images. Circularity (4π*(surface area/perimeter^2^)) represent the sphericity of mitochondria, value of 1 indicates perfect spheroids. Aspect ratio (AR) ((major axis)/(minor axis) reflects the ‘length-to-width ratio’. High aspect ratio represents better capacity of mitochondria to network.

### Seahorse assay

Extracellular acidification rate (ECAR) and oxygen consumption rate (OCR) analyses were performed with an XF96 extracellular flux analyzer (Seahorse Biosciences, North Billerica, MA) as described previously [48]. Briefly, 18×10^3^ cells were seeded per well on the XF96 cell plate overnight before the assay. OCR were measured in XF assay medium supplemented with 10 mM glucose, 2 mM glutamine and 1 mM sodium pyruvate followed by sequential addition of (i) 1 μM oligomycin; (ii) 0.5 μM FCCP, and (iii) 0.5 μM antimycin A + rotenone. ECAR were measured in XF assay medium with 0 mM glucose followed by (i) 10 mM glucose; (ii) 1 μM oligomycin; (iii) 50 mM 2-deoxyglucose (2-DG). Parameters of mitochondrial respiration were calculated as follows: basal respiration: (last measurement before oligomycin injection)-(non-mitochondrial oxygen consumption); maximal respiration: (maximum rate measurement after FCCP injection)-(non-mitochondrial oxygen consumption); spare respiratory capacity: (maximum respiration)-(basal respiration). The glycolysis ECAR was calculated as (ECAR after glucose addition)-(ECAR baseline). Results were normalized by protein content.

### SCENITH

SCENITH was performed as described in [31]. SCENITH™ reagents kit (inhibitors, puromycin and anti-Puromycin antibody clone R4743L-E8) was kindly provided by Dr Arguello and used according to protocol for in vitro cancer cell lines. Briefly, ovarian cancer cells were plated at 3.10^4^ cells/mL, in 24-well plates. Experimental triplicates were performed in all conditions. Cells were treated during 30min with Control, 2-Deoxy-D-Glucose (DG, 100mM), Oligomycin (Oligo, 1 μM), or the translation initiation inhibitor Harringtonine (2 μg/mL). Puromycin (10 μg/mL) is added during the last 15 min of the metabolic inhibitors treatment. After puromycin treatment, cells were detached and stained with Live/Dead reagent (ThermoFisher). Intracellular staining of puromycin was performed by using in house produced fluorescently labeled anti-puromycin monoclonal antibody with Alexa Fluor 647 by incubating cells during 1 hr at 4°C. Data acquisition was performed using the CYTOFLEX (Beckman Coulter) flow cytometer.

### Mitochondrial membrane potential measurements

Cells were incubated for 30 min at 37°C in a solution of modified Krebs Ringer buffer (mKRB: 135mM NaCl, 5mM KCl, 0.4mM KH_2_PO_4_, 1mM MgSO_4_, 20mM HEPES, 5.5mM glucose and 1mM CaCl_2_ (pH 7.4)) containing 2nM tetramethyl rhodamine methyl ester (TMRM; Life Technologies, T-668), Verapamil hydrochloride 20μM (Merck KGaA, V4629) and Hoechst 33342 1.6μM (Thermo Fisher, H3570).

Images were taken on a confocal laser scanning microscope (Olympus FLUOVIEW FV3000). TMRM excitation was performed at 561nm and emission was collected through a 570 to 620nm scanner.

TMRM average fluorescent units (AFU) were quantified through pixel densitometry quantification using ImageJ software. For each experimental condition, TMRM AFU were normalized for the control condition.

### Western Blot

For immunoblotting, cells were scraped into ice-cold, phosphate-buffered saline (PBS) and lysed in a modified 10 mM Tris buffer pH 7.4 containing 150 mM NaCl, 1% Triton X-100, 10% glycerol, 10 mM EDTA and protease and phosphatase inhibitor cocktail. After 30 min of incubation on ice, the lysates were cleared via centrifugation centrifuged at 12000 g at 4°C for 10 min. Samples protein concentration was assessed using the RC DC™ Protein Assay (Bio-Rad Laboratories). A volume correspondent to 20µg of protein was added to 1 volume of 4x Laemmli SDS sample loading buffer (Bio-Rad Laboratories), 1 volume of 10x NuPAGE™ sample reducing agent (Thermo Fisher, NP0009) and boiled 5 min at 100°C.

Samples were separated on a 4–12% Bolt™ Bis-Tris Plus Gels (Invitrogen, NW04127BOX) and electron-transferred to a nitrocellulose membrane for 3 hrs at 300 mA. Unspecific binding sites were saturated by incubating membranes with TBS-Tween 20 (0.05%) supplemented with 5% non-fat powdered milk for 1 hr at room temperature. Primary antibodies anti-Syntabulin (Genetex, GTX119953), anti-Mitofillin (Abcam, ab137057), total OXPHOS Human WB Antibody Cocktail (Abcam, ab110411), anti-GAPDH (Cell signalling technology, cs2118), anti-tubulin (Sigma, T9026) was performed over night at 4°C.

The next day, the revelation was assessed by the horseradish peroxidase (HRP) – conjugated secondary antibody (goat anti-rabbit, Thermo Fisher, G-21234) and the chemiluminescent substrate (Clarity Western ECL Substrate, Bio-Rad). Signal intensity was detected using FUSION FX digital imaging system (VILBER) and data analysis was performed via pixel densitometric quantification through ImageJ software.

### Metabolomic analysis

#### Sample preparation of cells and supernatants

COV318 cells were transfected for 48 hrs with control-siRNA or specific *SYBU* siRNA then lysed using cold methanol/water (9/1, v/v, −20°C) with internal standards (ISTD). Supernatants in microtubes were vortexed for 2500rpm for 5 min and centrifuged at 15000 g for 10 min at +4 °C. Supernatants were split in two parts: 150 µl were used for GC-MS experiment in injection vial and 300 µl were used for UHPLC-MS experimentation. After evaporation, dry GC-MS aliquot was spiked with 50 µl of methoxyamine (20 mg/ml in pyridine), stored at room temperature in dark overnight. Then, 80 µl of MSTFA was added and final derivatization occurred at 40°C during 30 minutes. Samples were directly injected into GC-MS. Regarding the LC-MS aliquots, aliquots were evaporated. The LC-MS dried extracts were solubilized with 150 µl of MilliQ water. Biological samples were kept at −80°C until injection or transferred in vials for direct analysis by UHPLC/MS.

### Widely-targeted analysis of intracellular metabolites gas chromatography (GC) coupled to a triple quadrupole (QQQ) mass spectrometer

GC-MS/MS method was performed on a coupling gas chromatography / triple quadrupole 7890B / 7000C (Agilent Technologies, Waldbronn, Germany). The scan mode used was the MRM for biological samples. Peak detection and integration of analytes were performed using the Agilent Mass Hunter quantitative software (B.07.01), exported as tables and processed with R software (version 4.0.3) and the GRMeta package (Github/kroemerlab) [58].

### Targeted analysis of nucleotides and cofactors by ion pairing ultra-high performance liquid chromatography (UHPLC) coupled to a Triple Quadrupole (QQQ) mass spectrometer

Targeted analysis was performed on a RRLC 1290 system (Agilent Technologies, Waldbronn, Germany) coupled to a Triple Quadrupole 6470 (Agilent Technologies) equipped with an electrospray source. Ten microliters of sample were injected on a Column Zorbax Eclipse XDB-C18 (100 mm x 2.1 mm particle size 1.8 µm) from Agilent technologies. Gradient mobile phase consisted of water with 2mM of dibutylamine acetate concentrate (DBAA) (A) and acetonitrile (B). Flow rate was set to 0.4 mL/min, and gradient as follow: initial condition was 90% phase A and 20% phase B, maintained during 3 min. Molecules were then eluted using a gradient from 10% to 95% phase B over 1 min. Column was washed using 95% mobile phase B for 2 minutes and equilibrated using 10% mobile phase B for 1 min. Scan mode used was the MRM for biological samples. Peak detection and integration of the analytes were performed using the Agilent Mass Hunter quantitative software (B.10.1).

MetaboAnalyst platform [59,60] (http://www.metaboanalyst.ca) was used to characterize major metabolic pathways regulated in cancer cells.

## Data availability

GR cohort transcriptomic data were deposited in the Biostudies ArrayExpress database under accession number E-MTAB-11177.

Mass spectrometry proteomics data have been deposited to the ProteomeXchange Consortium (http://proteomecentral.proteomexchange.org) via the PRIDE partner repository [61] with the dataset identifier PXD029075.

Raw metabolomic data from large-scale metabolomic analysis are publicly available in the Mendeley database (https://data.mendeley.com/datasets/2jtzpf3pjt/1).

Genomic data comparisons of *SYBU* expressions between human normal and tumoral tissues were obtained from the TNM plot database [60]. Kaplan–Meier HGSOC survival analysis was conducted from Kaplan–Meier Plotter (https://kmplot.com), for evaluating the patient prognosis according to their correlation with *SYBU* gene expression.

## Acknowledgements

Authors wish to thank Tudor Manoliu from the Imaging and Cytometry (PFIC) core facility, Bastien Job from the Bioinformatics platform and Sylvère Durand, Fanny Aprahamian and the metabolomics platform (INSERM US23 / CNRS UAR 3655, AMMICa of Université Paris-Saclay, Gustave Roussy Institute, Villejuif, France) for excellent expertise. We thank Léa Alban for valuable contribution to the project. The LSMP thanks Patrick Poullet from the bioinformatics Core facility (CUBIC) of the Institut Curie U1331 for the continuous development of myProMS.

