## Supplemental figures and legends for "Connecting mitochondrial metabolism and mitotic fidelity to control vulnerability of high grade serous ovarian cancer patients to taxane-based chemotherapy"

Hadia Moindjie et al.

### Supplementary Materials

#### Cell Lines

The human epithelial ovarian cancer cell lines COV318, OVCAR8, OCAR3, OVCAR4, OV56 and OV90 were kindly provided by Dr Fatima Mechta-Grigoriou (Institut Curie, Paris, France) and were cultured in Dulbecco's modified Eagle's medium (DMEM, GIBCO, Thermo Fisher Scientific) containing 10% fetal bovine serum (FBS, GIBCO, Thermo Fisher Scientific).

### Legends to Supplementary Figures

#### Supplementary Figure S1

A. Kaplan-Meier curves of progression-free survival analysis of HGSOC tumors classified as sensitive (S) or resistant (R) to taxane-based chemotherapy in the AOCS (left) and CURIE (right) cohorts.
B. Scattered dot plot of the SYBU probeset (218692_at) intensity in tumors from patients of the CURIE cohort classified as resistant (R) or sensitive (S) to neoadjuvant chemotherapy. Median values are highlighted in red.
C. Heatmap and hierarchical clustering of 42 HGSOC samples based on the intensities of SYBU probeset (218692_at) in the CURIE cohort. Left panel: Heatmap illustrating relative expression profiles of SYBU (column) for each tumor sample (line) in a continuous color scale from low (green) to high (red) expression. A dendrogram of the two selected tumor groups is shown on the right side. Right panel: Scattered dot plot of SYBU expression in each of the two selected clusters based on the dendrogram.
D. Percentage of tumors that are sensitive to chemotherapy among HGSOC in the CURIE cohort expressing high or low SYBU levels according to heatmap clustering (C).
E. Percentage of tumors with low SYBU levels in resistant (R) and sensitive (S) HGSOC groups in the CURIE cohort.
F. Scattered dot plot of SYBU probeset (218692_at) intensities in four major types of ovarian tumors in the CURIE cohort.

B-F. Number of tumors in each group is indicated in brackets.

* $P < 0.05$; **$P < 0.01$.

#### Supplementary Figure S2

A. Validation of SYBU depletion by real-time PCR following transfection of HeLa cells with SYBU-specific siRNA (siSYBU) for 72 hrs as compared to control (siCTR).
B. Real-time PCR analysis of SYBU mRNA levels in ovarian cancer cell lines.
C. Validation of SYBU depletion following transfection with two specific siRNA (siSYBU and siSYBU#2) in COV318 cells as in (A).
D. Representative immunoblot validating SYBU depletion by specific siRNA (si

SYBU) in COV318 cells. Blots were probed with anti-SYBU antibodies and anti-tubulin as internal control of equal protein loading.

E. Validation of SYBU depletion in COV318 as in (A) following permanent depletion of SYBU by specific shRNA (shSYBU and shSYBU#2) compared to control (shCTR) shRNA.
F. Representative immunoblot validating SYBU depletion by specific shRNA (shSYBU) in COV318 cells. Blots were probed as in (D).

*P* < 0.05; **P* < 0.01; ****P* < 0.0001.

**Supplementary Figure S3**

A. MetaboAnalyst 5.0 pathway enrichment analysis of metabolites significantly altered between COV318 cells expressing (siCTR) or not (siSYBU) SYBU (left) and in SYBU-depleted COV318 cells treated or not with PTX (5nM for 72 hrs) (right).
B. Quantification of the normalized relative abundance of 15 significantly altered metabolites (p-value <0,05) in the metabolomic analysis.

*P* < 0.05, **P* < 0.01

**Supplementary Figure S4**

A. Representative curves of oxygen consumption rates (OCR) in COV318 cells expressing (siCTR) or not (siSYBU or siSYBU#2) endogenous SYBU. O: oligomycin, F: FCCP, and R/A: rotenone/antimycin were sequentially injected to assess mitochondrial respiratory states.
B. Quantification of basal (left panel), maximal respiration (middle panel) and spare respiratory capacity (right panel) OCR in COV318 cells as in (A).
C. Representative curves of oxygen consumption rates (OCR) as in (A) in COV318 cells permanently silenced (shSYBU or shSYBU#2) or not (shCTR) for endogenous SYBU.
D. Quantification of basal (left panel), maximal respiration (middle panel) and spare respiratory capacity (right panel) OCR in COV318 cells as in (C).
E. Representative curves of oxygen consumption rates (OCR) as in (A) in OVCAR8 cells expressing SYBU (shCTR) or not (shSYBU).
F. Quantification of basal and maximal (left panel) and spare respiratory capacity (right panel) OCR in OVCAR8 cells as in (E).
G. Quantification of mitochondrial mass using mitotracker fluorescence intensity.

H. Quantification of mitochondrial DNA content by quantitative real time PCR.
I. Representative immunoblot showing abundance of 4 mitochondrial proteins (ATP5, UQCR2, SDHB and COXII) belonging to respiratory chain complexes (V, III, II AND IV) respectively, in cells expressing (siSYBU -) or not (siSYBU +) SYBU, treated with increasing doses of PTX as indicated. Vinculin immunostaining was the control of equal protein loading.
J. Quantification of TMRM fluorescence intensity (a.u : arbitrary unit) in COV318 cells permanently silenced (shSYBU) or not (shCTR) for SYBU following transfection with specific shRNA.
K. Analysis of mitochondrial dependence in COV318 cells expressing SYBU (siCTR) or not (siSYBU) by SCENITH™.
L. Quantification of extracellular acidification rate (ECAR) in COV318 cells expressing SYBU (siCTR) or not (siSYBU) SYBU following transfection with specific siRNA for 72 hrs.
M. Analysis of glycolytic capacity in COV318 cells expressing SYBU (siCTR) or not (siSYBU) by SCENITH™.
   $^{*}P < 0.05$, $^{**}P < 0.01$

**Supplementary Figure S5**

A-D. Quantification of mitochondrial surface (A), perimeter (B), circularity (C), and aspect ratio (D) from TEM images in Fig. 4 showing COV318 cells expressing (siCTR) or not (siSYBU) SYBU and treated or not with 5nM PTX during 72 hrs. The number of analyzed mitochondria is in brackets. $^{**}P < 0.01$

**Supplementary Figure S6**

A. Proteomaps-based GO enrichment analysis. Cell component (CC) treemap indicate enrichment significance based on adjusted P-value.
B. GO enrichment analysis. Biological process (BP) dotplot. BP column colours indicate enrichment significance based on adjusted P-value.
C. Representative images of in situ proximity ligation assay (PLA) analysis to control the specificity of PLA signals between GFP-SYBU and VDAC1, using mouse anti-GFP or rabbit anti-VDAC1 primary antibodies alone.
D. Cellular localization of GFP-SYBU and deletion mutants in COV318 cells determined by immunofluorescence using anti-GFP, anti-TOM20 (mitochondria) and anti-alpha-tubulin (microtubules) immunostaining.

**Supplementary Figure S7**

A. Representative images of in situ proximity ligation assay (PLA) analysis to control the specific interaction between MIC60 and VDAC in COV318 cells using either both mouse anti-MIC60 and rabbit anti-VDAC (left), or mouse anti-MIC60 primary antibodies only (right).

B. Representative images and quantification of PLA intensity signals per region of interest from (A). Number of cells is in brackets.

C-F. Quantification of mitochondrial surface (C), perimeter (D), circularity (E) and aspect ratio (F) in COV318 cells expressing (siCTR) or not (siSYBU) endogenous SYBU and transfected with GFP, GFP-SYBU and GFP-MIC60 as in Fig. 6B. Number of mitochondria is in brackets.

**$P$ < 0.01, ***$P$ < 0.005, ****$P$ < 0.0001.

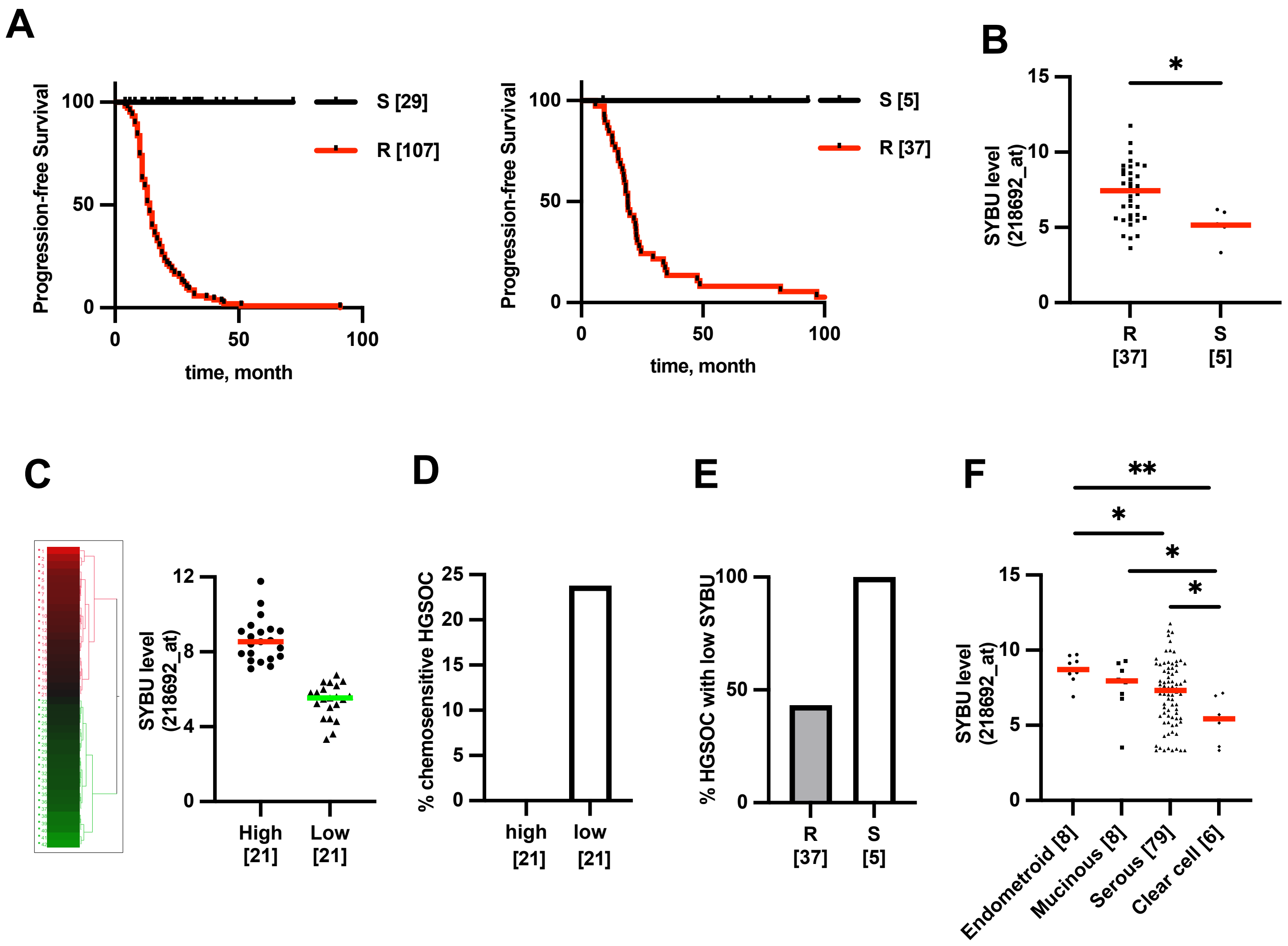

Moindjie et al., Suppl Figure S1

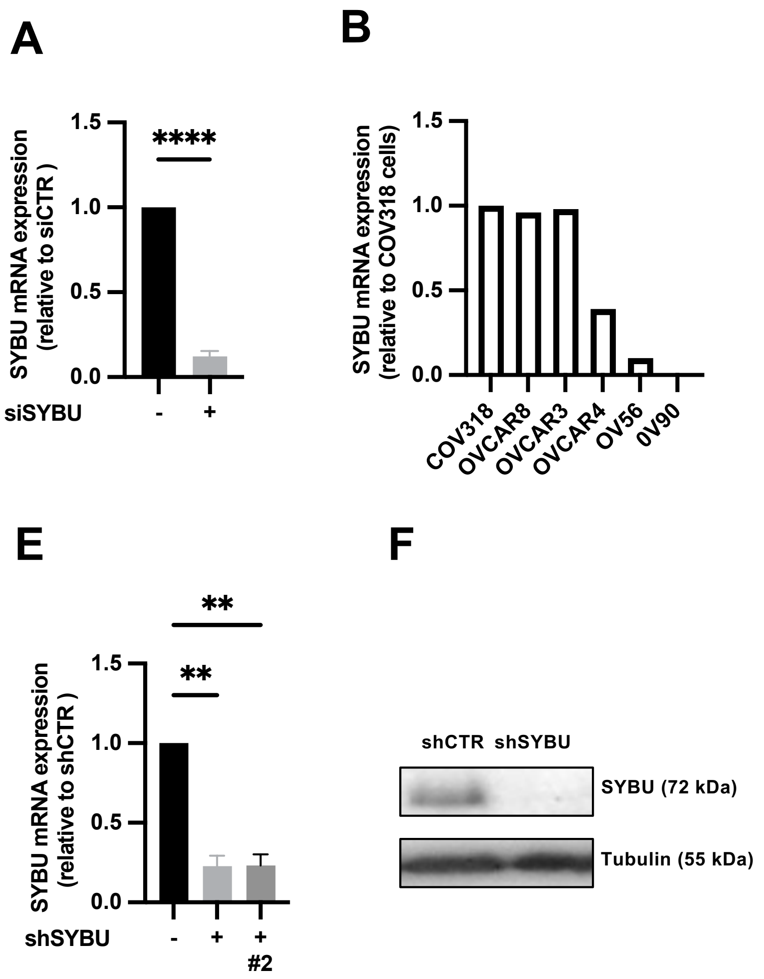

Moindjie et al., Suppl Figure S2

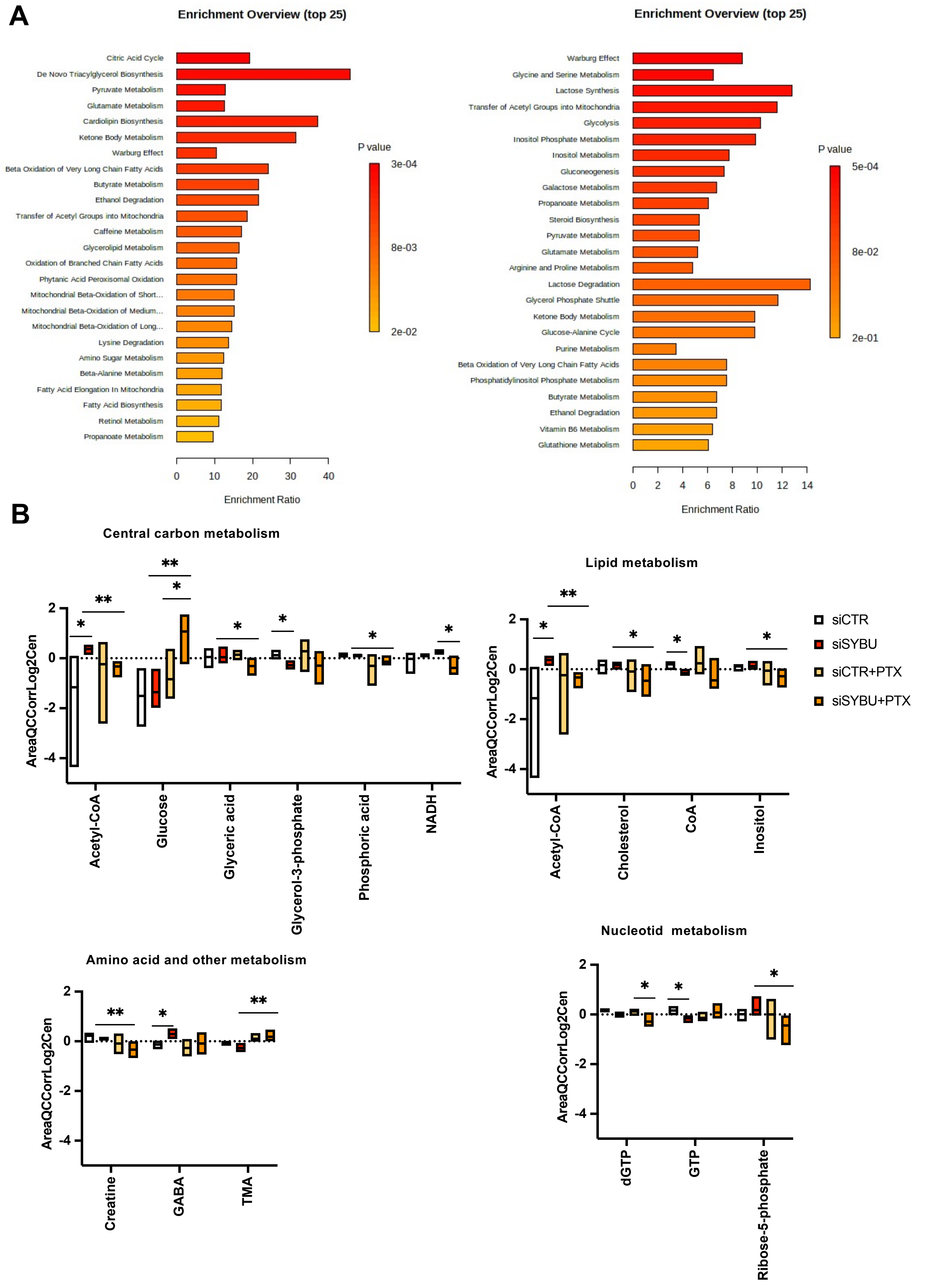

Moindjie et al., Suppl Figure S3

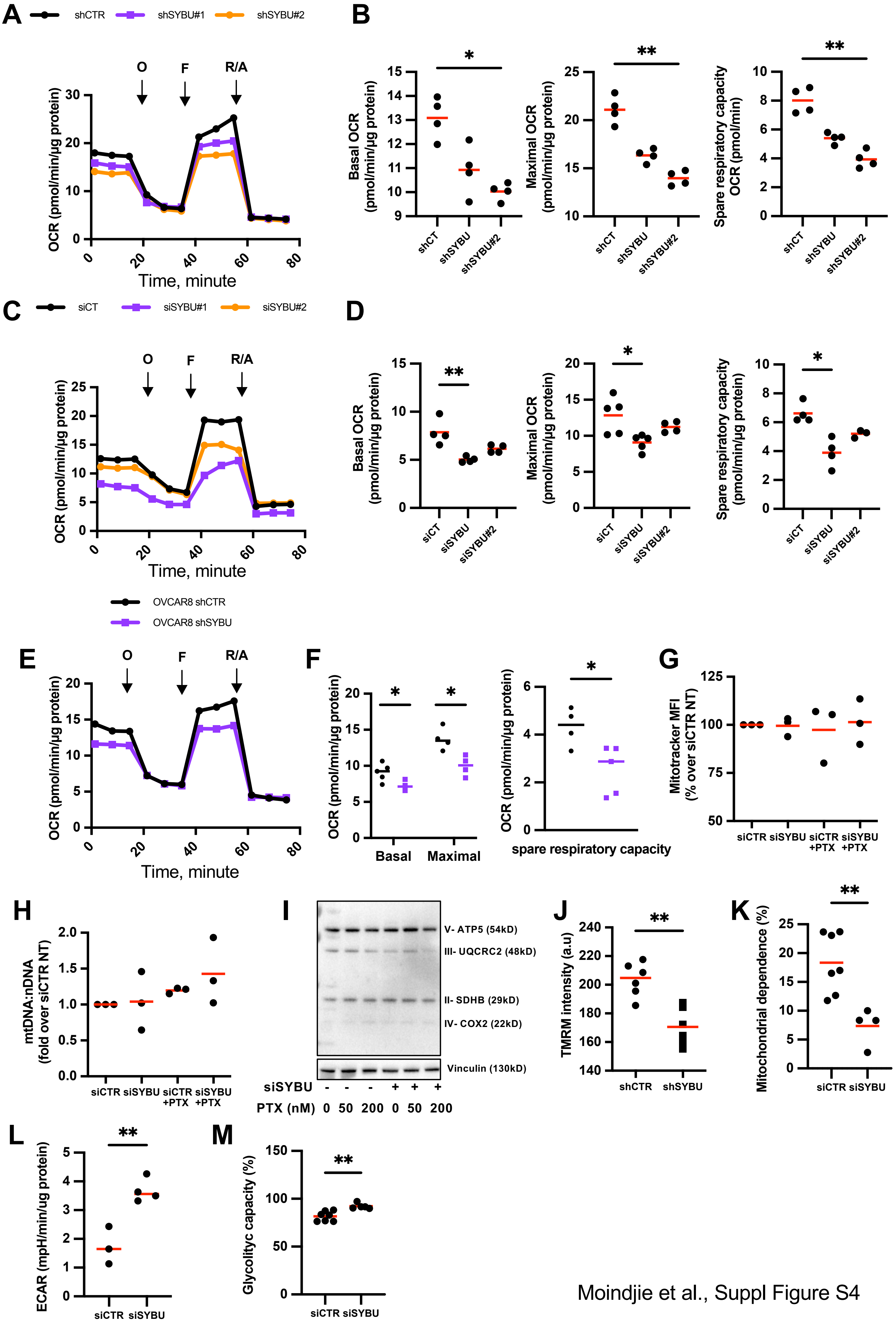

Moindjie et al., Suppl Figure S4

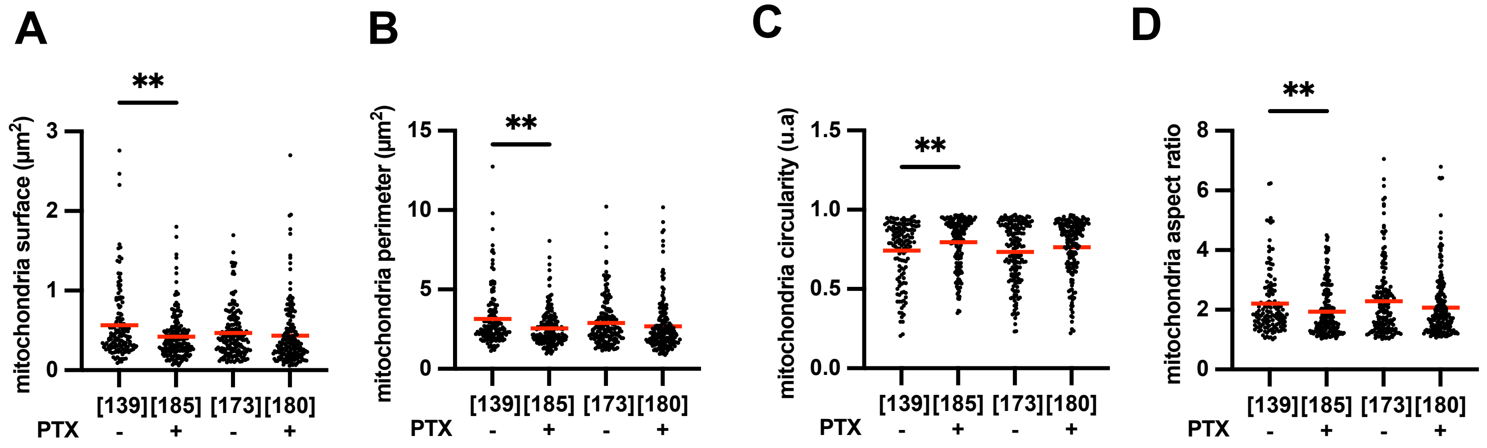

Moindjie et al., Suppl Figure S5

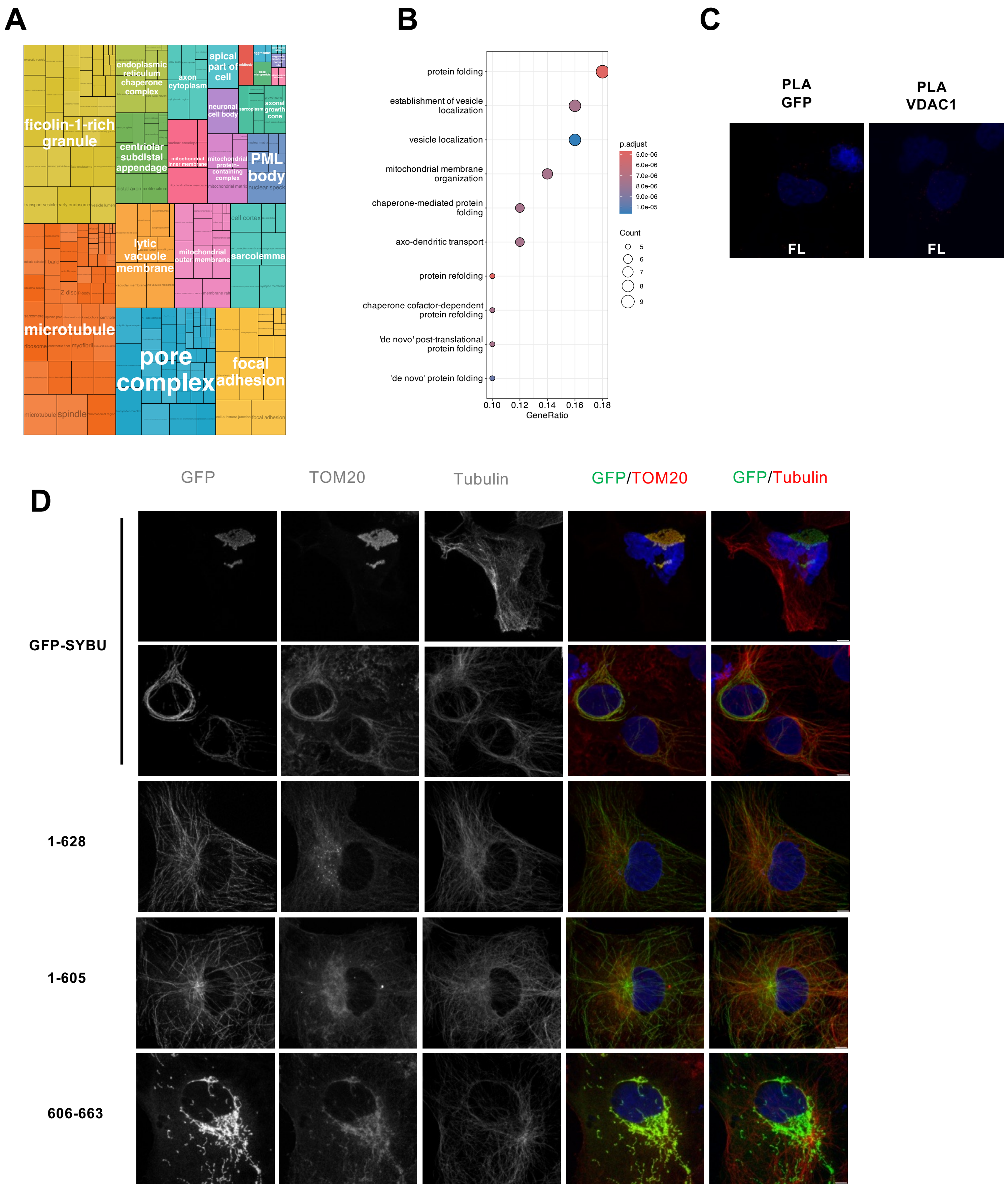

Moindjie et al., Suppl Figure S6

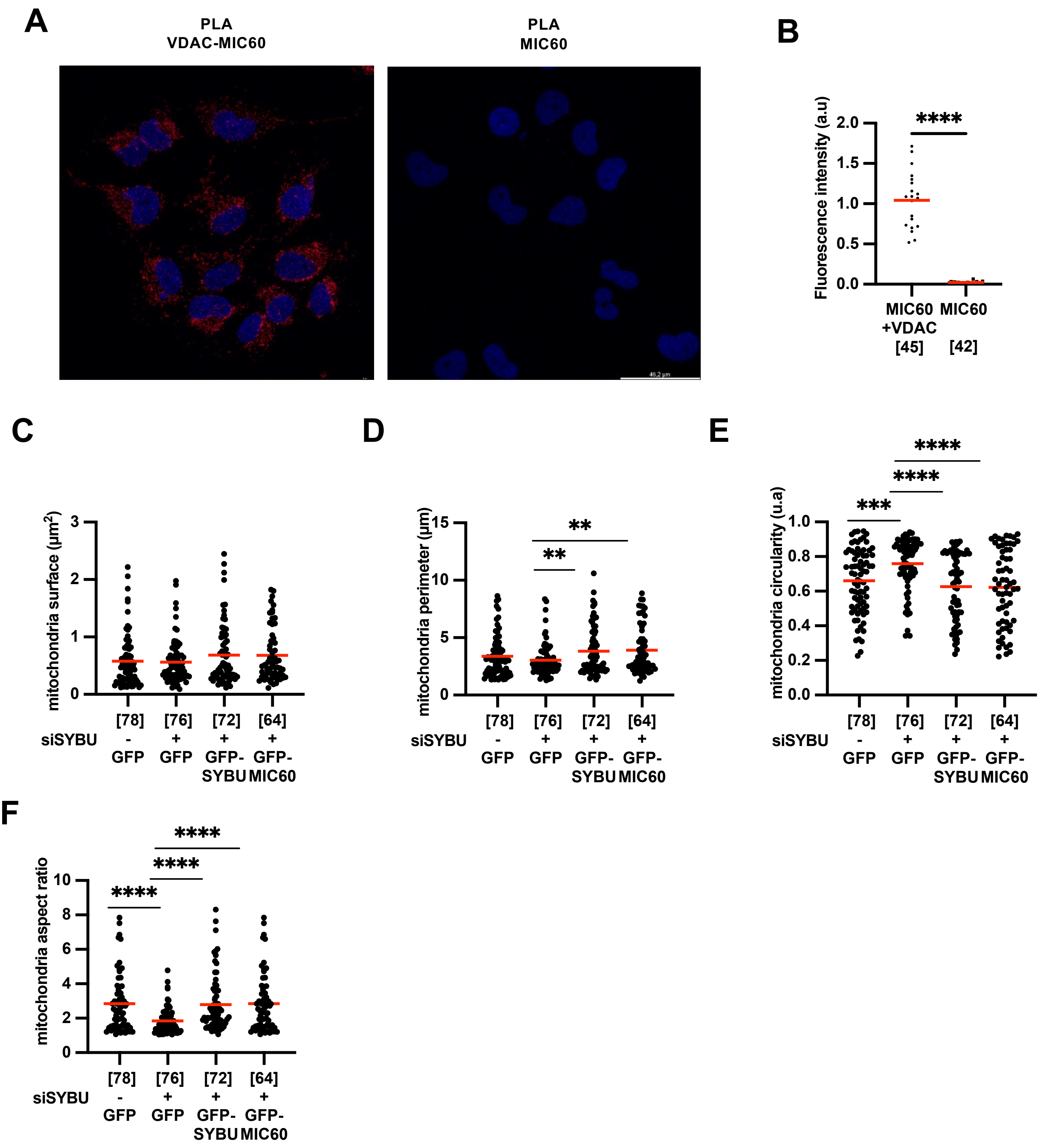

Moindjie et al., Suppl Figure S7

**Table S1. List of 410 microtubule-regulatory genes**

| Symbol | Gene Name | Gene ID |
|---|---|---|
| **AGTPBP1** | ATP/GTP binding protein 1 | 23287 |
| **AJUBA** | AJUBA LIM protein | 84962 |
| **AMER2** | APC membrane recruitment protein 2 | 219287 |
| **APC** | adenomatous polyposis coli | 324 |
| **APC2** | adenomatosis polyposis coli 2 | 10297 |
| **AR** | androgen receptor | 367 |
| **ARHGAP21** | Rho GTPase activating protein 21 | 57584 |
| **ARHGEF7** | Rho guanine nucleotide exchange factor (GEF) 7 | 8874 |
| **ASPM** | abnormal spindle microtubule assembly | 259266 |
| **ATAT1** | alpha tubulin acetyltransferase 1 | 79969 |
| **AURKA** | aurora kinase A | 6790 |
| **AURKB** | aurora kinase B | 9212 |
| **BAG3** | BCL2-associated athanogene 3 | 9531 |
| **BICD2** | BICD cargo adaptor 2 | 23299 |
| **BIRC5** | baculoviral IAP repeat containing 5 | 332 |
| **BOD1** | FAM44B | 91272 |
| **BRCA1** | breast cancer 1 | 672 |
| **BRCA2** | breast cancer 2 | 675 |
| **CAB39** | calcium binding protein 39 | 51719 |
| **CCND1** | cyclin-D1 | 595 |
| **CCT2** | chaperonin containing TCP1, subunit 2 (beta) | 10576 |
| **CDK1** | cyclin-dependent kinase 1 | 983 |
| **CDK2** | cyclin-dependent kinase 2 | 1017 |
| **CDK3** | cyclin-dependent kinase 3 | 1018 |
| **CDK4** | cyclin-dependent kinase 4 | 1019 |
| **CDK5** | cyclin-dependent kinase 5 | 1020 |
| **CDK5RAP2** | CDK5 regulatory subunit associated protein 2 | 55755 |
| **CDK6** | cyclin-dependent kinase 6 | 1021 |
| **CENPA** | centromere protein A | 1058 |
| **CENPB** | centromere protein B | 1059 |
| **CENPC** | centromere protein C | 1060 |
| **CENPF** | centromere protein F | 1063 |
| **CENPH** | centromere protein H | 64946 |
| **CENPJ** | centromere protein J | 55835 |
| **CKAP5** | CH-TOG | 9793 |
| **CLASP1** | cytoplasmic linker associated protein 1 | 23332 |
| **CLASP2** | cytoplasmic linker associated protein 2 | 23122 |
| **CLIP1** | CAP-GLY domain containing linker protein 1 | 6249 |
| **CLIP2** | CAP-GLY domain containing linker protein 2 | 7461 |
| **CLIP4** | CAP-GLY domain containing linker protein 4 | 79745 |
| **COPS5** | COP9 signalosome subunit 5 | 10987 |
| **COPS7A** | COP9 signalosome subunit 7A | 50813 |
| **COPS8** | COP9 signalosome subunit 8 | 10920 |
| **CSN3** | casein kappa | 1448 |
| **CSNK1D** | casein kinase 1, delta | 1453 |
| **CSNK1E** | casein kinase 1, epsilon | 1454 |
| **CSNK2A1** | casein kinase 2, alpha1 | 1457 |
| **CTNNB1** | catenin (cadherin-associated protein), beta 1, 88kDa | 1499 |
| **CTTNBP2** | Cortactin-binding protein 2 (CortBP2) | 83992 |
| **CUL7** | cullin 7 | 9820 |
| **CYFIP1** | cytoplasmic FMR1 interacting protein 1 | 23191 |
| **CYFIP2** | cytoplasmic FMR1 interacting protein 2 | 26999 |
| **CYLD** | cylindromatosis (turban tumor syndrome) | 1540 |
| **CYTSB** | SPECC1 sperm antigen with calponin homology and coiled-coil domains | 92521 |
| **DAW1** | dynein assembly factor with WD repeats 1 | 164781 |
| **DCDC2** | doublecortin domain containing 2 | 51473 |
| **DCTN1** | dynactin 1 (p150) | 1639 |
| **DCTN2** | dynactin 2 (p50) | 10540 |
| **DCX** | doublecortin | 1641 |
| **DDA3** | proline/serine-rich coiled-coil 1 (PSRC1) | 84722 |
| **DIAPH1** | diaphanous-related formin 1 | 1729 |
| **DIAPH2** | diaphanous-related formin 2 | 1730 |
| **DIAPH3** | diaphanous-related formin 3 | 81624 |
| **DISC1** | disrupted in schizophrenia 1 | 27185 |
| **DNAAF1** | dynein axonemal assembly factor 1 | 123872 |
| **DNAAF2** | dynein axonemal assembly factor 2 | 55172 |
| **DNAAF3** | dynein axonemal assembly factor 3 | 352909 |
| **DNAAF5** | dynein axonemal assembly factor 5 | 54919 |
| **DNAH1** | dynein axonemal heavy chain 1 | 25981 |
| **DNAH10** | dynein axonemal heavy chain 10 | 196385 |
| **DNAH11** | dynein axonemal heavy chain 11 | 8701 |
| **DNAH12** | dynein axonemal heavy chain 12 | 201625 |
| **DNAH14** | dynein axonemal heavy chain 14 | 127602 |
| **DNAH17** | dynein axonemal heavy chain 17 | 8632 |
| **DNAH2** | dynein axonemal heavy chain 2 | 146754 |
| **DNAH3** | dynein axonemal heavy chain 3 | 55567 |
| **DNAH5** | dynein axonemal heavy chain 5 | 1767 |
| **DNAH6** | dynein axonemal heavy chain 6 | 1768 |
| **DNAH7** | dynein axonemal heavy chain 7 | 56171 |
| **DNAH8** | dynein axonemal heavy chain 8 | 1769 |
| **DNAH9** | dynein axonemal heavy chain 9 | 1770 |
| **DNAI2** | dynein axonemal intermediate chain 2 | 64446 |
| **DNAI1** | dynein axonemal intermediate chain 1 | 27019 |
| **DNAL1** | dynein axonemal light chain 1 | 83544 |
| **DNAL4** | dynein axonemal light chain 4 | 10126 |
| **DNALI1** | dynein axonemal light intermediate chain 1 | 7802 |
| **DNHD1** | dynein heavy chain domain 1 | 144132 |
| **DNMT3B** | DNA (cytosine-5-)-methyltransferase 3 beta | 1789 |
| **DOCK7** | dedicator of cytokinesis 7 | 85440 |
| **DOT1L** | DOT1-like histone H3K79 methyltransferase | 84444 |
| **DPYSL2** | dihydropyrimidinase-like 2 | 1808 |
| **DRC1** | dynein regulatory complex subunit 1 | 92749 |
| **DRC3** | dynein regulatory complex subunit 3 | 83450 |
| **DRC7** | dynein regulatory complex subunit 7 | 84229 |
| **DST** | dystonin | 667 |
| **DYNC1H1** | dynein cytoplasmic 1 heavy chain 1 | 1778 |
| **DYNC1I1** | dynein, cytoplasmic 1, intermediate chain 1 | 1780 |
| **DYNC1I2** | dynein cytoplasmic 1 intermediate chain 2 | 1781 |
| **DYNC1LI1** | dynein cytoplasmic 1 light intermediate chain 1 | 51143 |
| **DYNC1LI2** | dynein cytoplasmic 1 light intermediate chain 2 | 1783 |
| **DYNC2H1** | dynein cytoplasmic 2 heavy chain 1 | 79659 |
| **DYNC2LI1** | dynein cytoplasmic 2 light intermediate chain 1 | 51626 |
| **DYNLL1** | dynein light chain LC8-type 1 | 8655 |
| **DYNLL2** | dynein light chain LC8-type 2 | 140735 |
| **DYNLRB1** | dynein light chain roadblock-type 1 | 83658 |
| **DYNLRB2** | dynein light chain roadblock-type 2 | 83657 |
| **DYNLT1** | dynein light chain Tctex-type 1 | 6993 |
| **DYNLT3** | dynein light chain Tctex-type 3 | 6990 |
| **EIF2AK2** | eukaryotic translation initiation factor 2-alpha kinase 2 | 5610 |
| **EML1** | echinoderm microtubule associated protein like 1 | 2009 |
| **EML2** | echinoderm microtubule associated protein like 2 | 24139 |
| **EML3** | echinoderm microtubule associated protein like 3 | 256364 |
| **EML4** | echinoderm microtubule associated protein like 4 | 27436 |
| **EML5** | echinoderm microtubule associated protein like 5 | 161436 |
| **EML6** | echinoderm microtubule associated protein like 6 | 400954 |
| **EP300** | E1A binding protein p300 | 2033 |
| **EPB41** | erythrocyte membrane protein band 4.1 | 2035 |
| **ERG** | v-ets avian erythroblastosis virus E26 oncogene homolog | 2078 |
| **FAM190A** | coiled-coil serine rich protein 1 (CCSER1) | 401145 |
| **FAM190B** | coiled-coil serine rich protein 2 (CCSER2) | 54462 |
| **FBL** | fibrillarin | 2091 |
| **FBXL14** | F-box and leucine-rich repeat protein 14 | 144699 |
| **FGFR1OP** | FGFR1 oncogene partner | 11116 |
| **FLIP1** | TNIP2 (TNFAIP3 interacting protein 2) | 79155 |
| **FMR1** | fragile X mental retardation 1 | 2332 |
| **FN1** | fibronectin 1 | 2335 |
| **FNTA** | farnesyltransferase, CAAX box, alpha | 2339 |
| **FNTB** | farnesyltransferase, CAAX box, beta | 2342 |
| **FYN** | FYN proto-oncogene, Src family tyrosine kinase | 2534 |
| **GAS2L1** | growth arrest specific 2 like 1 | 10634 |
| **GAS7** | growth arrest specific 7 [ | 8522 |
| **GIT1** | G protein-coupled receptor kinase interacting ArfGAP 1 | 28964 |
| **GOLGA2** | golgin A2 | 2801 |
| **GOLGB1** | golgin B1 | 2804 |
| **GPAA1** | glycosylphosphatidylinositol anchor attachment 1 | 8733 |
| **GSK3B** | glycogen synthase kinase 3 beta | 2932 |

| | | |
|---|---|---|
| **GSTP1** | glutathione S-transferase pi 1 | 2950 |
| **GTSE1** | G2 and S-phase expressed 1 | 51512 |
| **GZMB** | zyme B (granzyme 2, cytotoxic T-lymphocyte-associated serine esteras | 3002 |
| **HAUS6** | HAUS augmin like complex subunit 6 | 54801 |
| **HDAC6** | histone deacetylase 6 | 10013 |
| **HERC5** | HECT and RLD domain containing E3 ubiquitin protein ligase 5 | 51191 |
| **HSPA8** | heat shock 70kDa protein 8 | 3312 |
| **HTT** | huntingtin | 3064 |
| **iASPP** | PPP1R13L (protein phosphatase 1 regulatory subunit 13 like) | 10848 |
| **ID1** | inhibitor of DNA binding 1, dominant negative helix-loop-helix protein | 3397 |
| **INCENP** | inner centromere protein | 3619 |
| **IQCA1** | IQ motif containing with AAA domain 1 | 79781 |
| **IQCB1** | IQ motif containing B1 | 9657 |
| **IQCD** | IQ motif containing D | 115811 |
| **IQCG** | IQ motif containing G | 84223 |
| **IQGAP1** | IQ motif containing GTPase activating protein 1 | 8826 |
| **JAKMIP1** | janus kinase and microtubule interacting protein 1 | 152789 |
| **JAKMIP2** | janus kinase and microtubule interacting protein 2 | 9832 |
| **JAKMIP3** | Janus kinase and microtubule interacting protein 3 | 282973 |
| **KATNA1** | katanin p60 (ATPase containing) subunit A 1 | 11104 |
| **KDM5B** | lysine (K)-specific demethylase 5B | 10765 |
| **KDM6A** | lysine (K)-specific demethylase 6A | 7403 |
| **KIAA1524** | KIAA1524 (CIP2A) | 57650 |
| **KIF10** | kinesin family member 10=CENP-E | 1062 |
| **KIF11** | kinesin family member 11 = EG5 | 3832 |
| **KIF12** | kinesin family member 12 | 113220 |
| **KIF13A** | kinesin family member 13A | 63971 |
| **KIF13B** | kinesin family member 13B | 23303 |
| **KIF14** | kinesin family member 14 | 8928 |
| **KIF15** | kinesin family member 15 | 56992 |
| **KIF16A** | STARD9/KIF16A | 57519 |
| **KIF16B** | kinesin family member 16B | 55614 |
| **KIF17** | kinesin family member 17 | 57576 |
| **KIF18A** | kinesin family member 18A | 81930 |
| **KIF18B** | kinesin family member 18B | 146909 |
| **KIF19** | kinesin family member 19 | 124602 |
| **KIF1A** | kinesin family member 1A | 547 |
| **KIF1B** | kinesin family member 1B | 23095 |
| **KIF1C** | kinesin family member 1C | 10749 |
| **KIF20A** | kinesin family member 20A | 10112 |
| **KIF20B** | kinesin family member 20B | 9585 |
| **KIF21A** | kinesin family member 21A | 55605 |
| **KIF21B** | kinesin family member 21B | 23046 |
| **KIF22** | kinesin family member 22 | 3835 |
| **KIF23** | kinesin family member 23 | 9493 |
| **KIF24** | kinesin family member 24 | 347240 |
| **KIF25** | kinesin family member 25 | 3834 |
| **KIF26A** | kinesin family member 26A | 26153 |
| **KIF26B** | kinesin family member 26B | 55083 |
| **KIF27** | kinesin family member 27 | 55582 |
| **KIF2A** | kinesin family member 2A | 3796 |
| **KIF2B** | kinesin family member 2B | 84643 |
| **KIF2C** | kinesin family member 2C | 11004 |
| **KIF3A** | kinesin family member 3A | 11127 |
| **KIF3B** | kinesin family member 3B | 9371 |
| **KIF3C** | kinesin family member 3C | 3797 |
| **KIF4A** | kinesin family member 4A | 24137 |
| **KIF4B** | kinesin family member 4B | 285643 |
| **KIF5A** | kinesin family member 5A | 3798 |
| **KIF5B** | kinesin family member 5B | 3799 |
| **KIF5C** | kinesin family member 5C | 3800 |
| **KIF6** | kinesin family member 6 | 221458 |
| **KIF7** | kinesin family member 7 | 374654 |
| **KIF9** | kinesin family member 9 | 64147 |
| **KIFAP3** | kinesin associated protein 3 | 22920 |
| **KIFC1** | kinesin family member C1 | 3833 |
| **KIFC2** | kinesin family member C2 | 90990 |
| **KIFC3** | kinesin family member C3 | 3801 |
| **LASP1** | LIM and SH3 protein 1 | 3927 |
| **LGALS1** | lectin, galactoside-binding, soluble, 1 | 3956 |
| **LIMK1** | LIM domain kinase 1 | 3984 |
| **LIMK2** | LIM domain kinase 2 | 3985 |
| **LOC105376502** | microtubule cross-linking factor 1-like | 105376502 |
| **LOC107987035** | tubulin beta chain-like | 107987035 |
| **LRRK2** | leucine-rich repeat kinase 2 | 120892 |
| **MACF1** | microtubule-actin crosslinking factor 1 (ACF7) | 23499 |
| **MAP10** | microtubule-associated protein 10 | 54627 |
| **MAP1A** | microtubule associated protein 1A | 4130 |
| **MAP1B** | microtubule associated protein 1B (=MAP5) | 4131 |
| **MAP1LC3A** | microtubule associated protein 1 light chain 3 alpha | 84557 |
| **MAP1LC3B** | microtubule associated protein 1 light chain 3 beta | 81631 |
| **MAP1LC3C** | microtubule associated protein 1 light chain 3 gamma | 440738 |
| **MAP1S** | microtubule associated protein 1S (MAP8) | 55201 |
| **MAP2** | microtubule associated protein 2 | 4133 |
| **MAP4** | microtubule-associated protein 4 | 4134 |
| **MAP6** | microtubule-associated protein 6 | 4135 |
| **MAP7** | microtubule-associated protein 7 | 9053 |
| **MAP7D1** | MAP7 domain containing 1 | 55700 |
| **MAP7D2** | MAP7 domain containing 2 | 256714 |
| **MAP7D3** | MAP7 domain containing 3 | 79649 |
| **MAP9** | microtubule-associated protein 9 | 79884 |
| **MAPK3** | mitogen-activated protein kinase 3 | 5595 |
| **MAPK8IP1** | mitogen-activated protein kinase 8 interacting protein 1 | 9479 |
| **MAPRE1** | microtubule-associated protein, RP/EB family, member 1 | 22919 |
| **MAPRE2** | microtubule-associated protein, RP/EB family, member 2 | 10982 |
| **MAPRE3** | microtubule-associated protein, RP/EB family, member 3 | 22924 |
| **MAPT** | microtubule-associated protein tau | 4137 |
| **MARK1** | MAP/microtubule affinity-regulating kinase 1 | 4139 |
| **MARK2** | MAP/microtubule affinity-regulating kinase 2 | 2011 |
| **MARK3** | MAP/microtubule affinity-regulating kinase 3 | 4140 |
| **MARK4** | MAP/microtubule affinity-regulating kinase 4 | 57787 |
| **MAST1** | microtubule associated serine/threonine kinase 1 | 22983 |
| **MAST2** | microtubule associated serine/threonine kinase 2 | 23139 |
| **MAST3** | microtubule associated serine/threonine kinase 3 | 23031 |
| **MAST4** | microtubule associated serine/threonine kinase family member 4 | 375449 |
| **MASTL** | microtubule associated serine/threonine kinase-like | 84930 |
| **MCC** | mutated in colorectal cancers | 4163 |
| **MEMO-1** | mediator of cell motility 1 | 51072 |
| **MGEA5** | meningioma expressed antigen 5 (hyaluronidase) | 10724 |
| **MICAL1** | rotubule associated monooxygenase, calponin and LIM domain containi | 64780 |
| **MICAL2** | rotubule associated monooxygenase, calponin and LIM domain containi | 9645 |
| **MICAL3** | rotubule associated monooxygenase, calponin and LIM domain containi | 57553 |
| **MICALCL** | MICAL C-terminal like | 84953 |
| **MICALL1** | MICAL-like 1 | 85377 |
| **MICALL2** | MICAL-like 2 | 79778 |
| **MITD1** | MIT, microtubule interacting and transport, domain containing 1 | 129531 |
| **MITF** | microphthalmia-associated transcription factor | 4286 |
| **MLPH** | melanophilin | 79083 |
| **MTCL1** | microtubule crosslinking factor 1 | 23255 |
| **MTOR** | mechanistic target of rapamycin | 2475 |
| **MTUS1** | microtubule associated tumor suppressor 1 | 57509 |
| **MTUS2** | microtubule associated tumor suppressor candidate 2 | 23281 |
| **MZT1** | mitotic spindle organizing protein 1 | 440145 |
| **MZT2A** | mitotic spindle organizing protein 2A | 653784 |
| **MZT2B** | mitotic spindle organizing protein 2B | 80097 |
| **NAV1** | neuron navigator 1 | 89796 |
| **NAV2** | neuron navigator 2 | 89797 |
| **NAV3** | neuron navigator 3 | 89795 |
| **NCK5L** | NCKAP5L (NCK associated protein 5 like) | 57701 |
| **NCKP5** | NCKAP5 (NCK associated protein 5) | 344148 |
| **NCOA6** | nuclear receptor coactivator 6 | 23054 |
| **NDEL1** | nudE neurodevelopment protein 1-like 1 | 81565 |
| **NEK1** | NIMA related kinase 1 | 4750 |
| **NEK10** | NIMA related kinase 10 | 152110 |
| **NEK11** | NIMA related kinase 11 | 79858 |
| **NEK2** | NIMA related kinase 2 | 4751 |
| **NEK3** | NIMA related kinase 3 | 4752 |
| **NEK4** | NIMA related kinase 4 | 6787 |
| **NEK5** | NIMA related kinase 4 | 341676 |
| **NEK6** | NIMA related kinase 6 | 10783 |
| **NEK7** | NIMA related kinase 7 | 140609 |

| | | |
|---|---|---|
| **NEK8** | NIMA related kinase 8 | 284086 |
| **NEK9** | NIMA related kinase 9 | 91754 |
| **NOTCH1** | notch 1 | 4851 |
| **NUDC** | nuclear distribution C, dynein complex regulator | 10726 |
| **NUMA1** | nuclear mitotic apparatus protein 1 | 4926 |
| **NUPR1** | nuclear protein 1, transcriptional regulator | 26471 |
| **PAK1** | p21 protein (Cdc42/Rac)-activated kinase 1 | 5058 |
| **PARD6G** | par-6 family cell polarity regulator gamma | 84552 |
| **PARK2** | parkin RBR E3 ubiquitin protein ligase | 5071 |
| **PAXIP1** | PAX interacting (with transcription-activation domain) protein 1 | 22976 |
| **PCNA** | proliferating cell nuclear antigen | 5111 |
| **PDLIM5** | PDZ and LIM domain 5 | 10611 |
| **PIN1** | peptidylprolyl cis/trans isomerase, NIMA-interacting 1 | 5300 |
| **PIN4** | peptidylprolyl cis/trans isomerase, NIMA-interacting 4 | 5303 |
| **PKN1** | protein kinase N1 | 5585 |
| **PLK1** | polo like kinase 1 | 5347 |
| **PLXNA2** | plexin A2 | 5362 |
| **PNMA2** | paraneoplastic Ma antigen 2 | 10687 |
| **PRKAA1** | protein kinase AMP-activated catalytic subunit alpha 1 | 5562 |
| **PRKAA2** | protein kinase AMP-activated catalytic subunit alpha 2 | 5563 |
| **PRMT1** | protein arginine methyltransferase 1 | 3276 |
| **PSEN1** | presenilin 1 | 5663 |
| **PSEN2** | presenilin 2 | 5664 |
| **PXN** | paxillin | 5829 |
| **RACGAP1** | Rac GTPase activating protein 1 | 29127 |
| **RAN** | RAN, member RAS oncogene family | 5901 |
| **RASSF1** | Ras association (RalGDS/AF-6) domain family member 1 | 11186 |
| **RB1** | retinoblastoma 1 | 5925 |
| **RBBP5** | retinoblastoma binding protein 5 | 5929 |
| **REST** | RE1-silencing transcription factor | 5978 |
| **RITA1** | RBPJ interacting and tubulin associated 1 | 84934 |
| **RMDN1** | regulator of microtubule dynamics 1 | 51115 |
| **RMDN2** | regulator of microtubule dynamics 2 | 151393 |
| **RMDN3** | regulator of microtubule dynamics 3 | 55177 |
| **RPTOR** | regulatory associated protein of MTOR, complex 1 | 57521 |
| **SATB1** | SATB homeobox 1 | 6304 |
| **SEL1L** | sel-1 suppressor of lin-12-like (C. elegans) | 6400 |
| **SFPQ** | splicing factor proline/glutamine-rich | 6421 |
| **SGOL1** | shugoshin-like 1 | 151648 |
| **SGOL2** | shugoshin-like 2 | 151246 |
| **SHOX2** | short stature homeobox 2 | 6474 |
| **SIAH1** | siah E3 ubiquitin protein ligase 1 | 6477 |
| **SIRT2** | sirtuin 2 | 22933 |
| **SKAP** | KNSTRN (kinetochore-localized astrin/SPAG5 binding protein) | 90417 |
| **SKI** | SKI proto-oncogene | 6497 |
| **SLAIN1** | SLAIN motif family member 1 | 122060 |
| **SLAIN2** | SLAIN motif family member 2 | 57606 |
| **SMARCA4** | ted, matrix associated, actin dependent regulator of chromatin, subfamily | 6597 |
| **SMC3** | structural maintenance of chromosomes 3 | 9126 |
| **SNCA** | synuclein alpha | 6622 |
| **SPAST** | spastin | 6683 |
| **SPI1** | Spi-1 proto-oncogene | 6688 |
| **SRCIN1** | SRC kinase signaling inhibitor 1 | 80725 |
| **SRF** | serum response factor | 6722 |
| **STAU1** | staufen double-stranded RNA binding protein 1 | 6780 |
| **STIM1** | stromal interaction molecule 1 | 6786 |
| **STMN1** | stathmin 1 | 3925 |
| **STRADA** | STE20-related kinase adaptor alpha | 92335 |
| **SUZ12** | SUZ12 polycomb repressive complex 2 subunit | 23512 |
| **SVIL** | supervillin | 6840 |
| **SYBU** | syntabulin | 55638 |
| **SYK** | spleen tyrosine kinase | 6850 |
| **Tastin** | TROAP (trophinin associated protein) | 10024 |
| **TBCA** | tubulin folding cofactor A | 6902 |
| **TBCB** | tubulin folding cofactor B | 1155 |
| **TBCC** | tubulin folding cofactor C | 6903 |
| **TBCD** | tubulin folding cofactor D | 6904 |
| **TBCE** | tubulin folding cofactor E | 6905 |
| **TBCEL** | tubulin folding cofactor E like | 219899 |
| **TERF1** | telomeric repeat binding factor (NIMA-interacting) 1 | 7013 |
| **TERF2** | telomeric repeat binding factor 2 | 7014 |
| **TGM2** | transglutaminase 2 | 7052 |
| **TP53** | tumor protein p53 | 7157 |
| **TPGS1** | tubulin polyglutamylase complex subunit 1 | 91978 |
| **TPGS2** | tubulin polyglutamylase complex subunit 2 | 25941 |
| **TPPP** | tubulin polymerization promoting protein | 11076 |
| **TPPP2** | tubulin polymerization promoting protein family member 2 | 122664 |
| **TPPP3** | tubulin polymerization promoting protein family member 3 | 51673 |
| **TPX2** | TPX2, microtubule-associated | 22974 |
| **TRAF6** | TNF receptor-associated factor 6, E3 ubiquitin protein ligase | 7189 |
| **TRIP10** | thyroid hormone receptor interactor 10 | 9322 |
| **TTBK1** | tau tubulin kinase 1 | 84630 |
| **TTBK2** | tau tubulin kinase 2 | 146057 |
| **TTL** | tubulin tyrosine ligase | 150465 |
| **TTLL1** | tubulin tyrosine ligase like 1 | 25809 |
| **TTLL10** | tubulin tyrosine ligase like 10 | 254173 |
| **TTLL11** | tubulin tyrosine ligase like 11 | 158135 |
| **TTLL12** | tubulin tyrosine ligase like 12 | 23170 |
| **TTLL2** | tubulin tyrosine ligase like 2 | 83887 |
| **TTLL3** | tubulin tyrosine ligase like 3 | 26140 |
| **TTLL4** | tubulin tyrosine ligase like 4 | 9654 |
| **TTLL5** | tubulin tyrosine ligase like 5 | 23093 |
| **TTLL6** | tubulin tyrosine ligase like 6 | 284076 |
| **TTLL7** | tubulin tyrosine ligase like 7 | 79739 |
| **TTLL8** | tubulin tyrosine ligase like 8 | 164714 |
| **TTLL9** | tubulin tyrosine ligase like 9 | 164395 |
| **TUBA1A** | tubulin, alpha 1a | 7846 |
| **TUBA1B** | tubulin, alpha 1b | 10376 |
| **TUBA1C** | tubulin, alpha 1c | 84790 |
| **TUBA3C** | tubulin, alpha 3c | 7278 |
| **TUBA3D** | tubulin, alpha 3d | 113457 |
| **TUBA3E** | tubulin, alpha 3e | 112714 |
| **TUBA4A** | tubulin, alpha 4a | 7277 |
| **TUBA4B** | tubulin, alpha 4b | 80086 |
| **TUBA8** | tubulin, alpha 8 | 51807 |
| **TUBAL3** | tubulin alpha like 3 | 79861 |
| **TUBB** | tubulin, beta class I | 203068 |
| **TUBB1** | tubulin, beta 1 class VI | 81027 |
| **TUBB2A** | tubulin, beta 2A class IIa | 7280 |
| **TUBB2B** | tubulin beta 2B class IIb | 347733 |
| **TUBB3** | tubulin, beta 3 class III | 10381 |
| **TUBB4A** | tubulin, beta 4A class IVa | 10382 |
| **TUBB4B** | tubulin, beta 4B class IVb | 10383 |
| **TUBB6** | tubulin, beta 6, class V | 84617 |
| **TUBB8** | tubulin, beta 8, class VIII | 347688 |
| **TUBD1** | tubulin, delta 1 | 51174 |
| **TUBE1** | tubulin, epsilon 1 | 51175 |
| **TUBG1** | tubulin, gamma 1 | 7283 |
| **TUBG2** | tubulin, gamma 2 | 27175 |
| **TUBGCP2** | tubulin, gamma complex associated protein 2 | 10844 |
| **TUBGCP3** | tubulin, gamma complex associated protein 3 | 10426 |
| **TUBGCP4** | tubulin, gamma complex associated protein 4 | 27229 |
| **TUBGCP5** | tubulin, gamma complex associated protein 5 | 114791 |
| **TUBGCP6** | tubulin, gamma complex associated protein 6 | 85378 |
| **UBC** | ubiquitin C | 7316 |
| **UCHL1** | ubiquitin C-terminal hydrolase L1 | 7345 |
| **USP21** | ubiquitin specific peptidase 21 | 27005 |
| **VHL** | von Hippel-Lindau tumor suppressor, E3 ubiquitin protein ligase | 7428 |
| **WASH1** | WAS protein family homolog 1 | 100287171 |
| **WDR5** | WD repeat domain 5 | 11091 |
| **WHAMM** | S protein homolog associated with actin, golgi membranes and microtub | 123720 |
| **WHSC1** | Wolf-Hirschhorn syndrome candidate 1 | 7468 |
| **ZBED6** | zinc finger, BED-type containing 6 | 100381270 |

**Table S2. Clinical characteristics of HGSOC cohorts of patients**

| | ***AOCS cohort*** n= 282<br>GSE 9899 | ***CURIE cohort*** n= 101<br>GSE 26193 |
|---|---|---|
| ***Median age (years)*** | 59 | 57,8 |
| Range | 22-80 | 31,1-86,8 |
| < 65 years | 191 | 73 |
| ≥ 65 years | 91 | 28 |
| ***Grade*** | | |
| 1 | 20 (7,1%) | 7 (6,9%) |
| 2 | 99 (35,1%) | 32 (31,7%) |
| 3 | 160 (56,7%) | 62 (61,4%) |
| Unknown | 3 (1,1%) | 0 (0%) |
| ***FIGO stage at diagnosis*** | | |
| I | 24 (8,5%) | 21 (20,8%) |
| II | 18 (6,4%) | 9 (8,9%) |
| III | 217 (77%) | 56 (55,4%) |
| IV | 22 (7,8) | 15 (14,9%) |
| Unknown | 1 (0,3%) | 0 |
| ***Chemotherapy*** | | |
| Yes | 243 (86,2%) | 87 (86,1%) |
| No | 39 (13,8%) | 14 (13,9%) |
| ***Relapse*** | | |
| Yes | 190 (67,4%) | 75 (74,3%) |
| No | 92 (32,6%) | 26 (25,7%) |
| ***Progression free-survival (months)*** | | |
| Median months | 15 | 19,4 |
| Range | 1-166 | 0,1-243 |
| < 12 months | 105 | 25 |
| > 12 months | 177 | 76 |
| ***Overall survival*** | | |
| Median months | 28 | 36,5 |
| Dead | 169 (60%) | 71 (70,3%) |
| Alive | 114 (40%) | 30 (29,7%) |

**Table S3: Microtubule-regulatory genes differentially expressed between sensitive (S) and resistant (R) HGSOC tumors from patients included in the AOCS (panel A) and the CURIE (panel B) cohorts.**

A

| AOCS | | | |
|---|---|---|---|
| **Symbol** | **ProbeSet** | **p-value** | **Fold-change S/R** |
| **CENPB** | 212437_at | 0.0129466 | 1.2 |
| **CENPJ** | 223513_at | 0.0097447 | 0.78 |
| **CLASP2** | 1558759_s_at | 0.0082431 | 1.26 |
| **CTNNB1** | 1554411_at | 0.000985 | 1.42 |
| **DNAH9** | 240857_at | 0.0043346 | 1.33 |
| **DNAL1** | 223959_at | 0.0060071 | 1.27 |
| **DYNLRB2** | 238116_at | 0.0110902 | 1.59 |
| **EML6** | 229656_s_at | 0.0226696 | 1.34 |
| **FBXL14** | 213145_at | 0.0422168 | 1.2 |
| **FNTB** | 1568865_at | 0.0257653 | 1.21 |
| **KDM6A** | 203990_s_at | 0.0294005 | 1.23 |
| **KIF3A** | 213623_at | 0.0111068 | 1.3 |
| **KIF6** | 1562639_at | 0.0005148 | 1.26 |
| **KIFC1** | 209680_s_at | 0.0206574 | 0.81 |
| **LIMK2** | 202193_at | 0.0461123 | 0.78 |
| **MICAL1** | 218376_s_at | 0.0196053 | 0.81 |
| **MICAL3** | 212715_s_at | 0.0082759 | 0.73 |
| **NEK4** | 204634_at | 0.011873 | 1.22 |
| **NUPR1** | 209230_s_at | 0.0304779 | 0.73 |
| **PNMA2** | 209598_at | 0.0402074 | 1.58 |
| **SIAH1** | 232365_at | 0.0197242 | 1.29 |
| **SNCA** | 204466_s_at | 0.0331189 | 0.75 |
| **SPAST** | 207724_s_at | 0.0048581 | 1.24 |
| **SYBU** | **218692_at** | **0.0228395** | **0.61** |
| **TUBGCP4** | 211337_s_at | 0.0031247 | 1.25 |

B

| CURIE | | | |
|---|---|---|---|
| **Symbol** | **Probe set** | **p-value** | **Fold-change S/R** |
| **ATAT1** | 228510_at | 0.0274815 | 1.99 |
| **CAB39** | 224311_s_at | 0.0355172 | 1.86 |
| | 217873_at | 0.039847 | 1.45 |
| **CYFIP2** | 220999_s_at | 0.0066414 | 2.6 |
| | 215785_s_at | 0.0472145 | 2.25 |
| **FN1** | 1558199_at | 0.0017436 | 2.72 |
| | 214701_s_at | 0.0062915 | 5.09 |
| | 212464_s_at | 0.0091731 | 3.81 |
| | 214702_at | 0.0102597 | 3.69 |
| | 210495_x_at | 0.0119155 | 3.64 |
| | 216442_x_at | 0.0158025 | 3.58 |
| | 211719_x_at | 0.0184839 | 3.42 |
| **KDM6A** | 203992_s_at | 0.0282172 | 1.54 |
| **KIF1B** | 209234_at | 6.22e-05 | 0.4 |
| **LASP1** | 200618_at | 0.013535 | 1.52 |
| **LRRK2** | 229584_at | 0.0096666 | 2.04 |
| **MAP1A** | 203151_at | 8,00E-06 | 2.56 |
| **MAP1B** | 214577_at | 0.0280867 | 2.14 |
| **MAPRE2** | 202501_at | 0.0223881 | 1.98 |
| **MARK3** | 202569_s_at | 0.0091903 | 1.69 |
| **MICAL1** | 218376_s_at | 0.0061389 | 1.81 |
| **NEK11** | 1555082_a_at | 0.0261379 | 0.6 |
| | 219542_at | 0.0268523 | 0.53 |
| **NEK9** | 212299_at | 0.0251024 | 0.53 |
| **PLXNA2** | 213030_s_at | 0.0207336 | 2.77 |
| **SYBU** | **218692_at** | **0.0215106** | **0.23** |
| **TTBK2** | 1557073_s_at | 0.0055636 | 1.95 |
| | 213922_at | 0.0159373 | 1.55 |
| **TUBG2** | 203894_at | 0.0244881 | 1.51 |
| **VHL** | 1559227_s_at | 0.048734 | 0.66 |

Table S4. MT-regulating genes differentially expressed between Borderline (BOL) and adenocarcinoma (ADK) tumors in IGR cohort

| IGR | | |
|---|---|---|
| Symbol | p-value | Fold-change ADK/BOL |
| ASPM | < 1e-07 | 2.72 |
| AURKA | < 1e-07 | 1.99 |
| BIRC5 | < 1e-07 | 4.39 |
| CCND1 | 6.1e-05 | 0.58 |
| CDK1 | < 1e-07 | 3.11 |
| CENPA | < 1e-07 | 2.65 |
| CENPF | < 1e-07 | 4.37 |
| CLIP4 | 9.81e-05 | 0.64 |
| CSNK1E | 1.91e-05 | 0.62 |
| DIAPH3 | < 1e-07 | 1.42 |
| DNAH12 | 1,00E-07 | 0.51 |
| DNAH2 | < 1e-07 | 0.45 |
| DNAH5 | < 1e-07 | 0.45 |
| DNAH7 | < 1e-07 | 0.65 |
| DNAH9 | < 1e-07 | 0.45 |
| DNAI2 | 3.8e-06 | 0.65 |
| DNALI1 | 2.01e-05 | 0.36 |
| DNMT3B | < 1e-07 | 2.04 |
| DPYSL2 | 0.0055654 | 0.63 |
| DST | 3.3e-06 | 0.65 |
| DYNC2H1 | 2,00E-07 | 0.7 |
| DYNLRB2 | < 1e-07 | 0.23 |
| DYNLT1 | 0.002121 | 0.7 |
| EIF2AK2 | 2,00E-07 | 1.58 |
| GSTP1 | 0.0088996 | 1.4 |
| HAUS6 | 8.4e-06 | 1.55 |
| IQCA1 | 0.0053841 | 0.64 |
| IQCD | 6.1e-06 | 0.49 |
| IQCG | 6.23e-05 | 0.63 |
| KIF11 | < 1e-07 | 1.79 |
| KIF13B | < 1e-07 | 0.55 |
| KIF18A | < 1e-07 | 1.43 |
| KIF18B | < 1e-07 | 1.84 |
| KIF20A | < 1e-07 | 1.53 |
| KIF2C | < 1e-07 | 1.67 |
| KIF9 | < 1e-07 | 0.42 |
| KIFC1 | < 1e-07 | 1.79 |
| MAP9 | < 1e-07 | 0.43 |
| MAPRE1 | 2,00E-07 | 1.53 |
| MAST4 | 2,00E-06 | 0.6 |
| MICALL2 | 4,00E-07 | 0.48 |
| MLPH | 8.84e-05 | 0.54 |
| MZT1 | 4,00E-07 | 1.48 |
| NEK2 | < 1e-07 | 1.99 |
| PCNA | 6,00E-07 | 1.74 |
| PDLIM5 | < 1e-07 | 0.59 |
| PLK1 | < 1e-07 | 1.61 |
| PSRC1 | < 1e-07 | 1.5 |
| RACGAP1 | < 1e-07 | 3.37 |
| RAN | 4,00E-07 | 1.54 |
| SLAIN1 | 3.81e-05 | 0.51 |
| STMN1 | < 1e-07 | 3.02 |
| SYBU | 2.8e-06 | 0.58 |
| TBCE | < 1e-07 | 1.62 |
| TPPP3 | < 1e-07 | 0.33 |
| TPX2 | < 1e-07 | 1.51 |
| TTLL12 | < 1e-07 | 0.6 |
| TUBA1B | < 1e-07 | 2.25 |
| TUBA1C | < 1e-07 | 1.76 |
| TUBB2A | 6.7e-06 | 1.51 |
| TUBB4B | 0.0025899 | 0.66 |
| UBC | 6.1e-06 | 1.52 |
| UCHL1 | 0.0001154 | 2.39 |
| VHL | < 1e-07 | 1.41 |

**Table S5: Differential metabolites detected in COV318 cells, pvalue and fold change value**

| | siSYBU/siCTR | | siCTR+PTX/siCTR | | siSYBU+PTX/siSYBU | | siSYBU+PTX/siCTR+PTX | |
|---|---|---|---|---|---|---|---|---|
| | p-value | Fold-change | pvalue | Fold-change | pvalue | Fold-change | pvalue | Fold-change |
| **Acetyl CoA** | **0,02** | **1,53** | 0,20 | 0,93 | **0,01** | **-0,70** | 0,29 | -0,09 |
| **Cholesterol** | 1,00 | 0,00 | 1,00 | -0,26 | **0,03** | **-0,62** | 0,29 | -0,37 |
| **CoA** | **0,03** | **-0,35** | 0,89 | 0,05 | 0,15 | -0,29 | 0,11 | -0,68 |
| **Creatine** | 0,29 | -0,10 | 0,20 | -0,31 | **0,01** | **-0,46** | 0,41 | -0,25 |
| **dGTP** | 0,06 | -0,15 | 0,89 | -0,03 | 0,06 | -0,28 | **0,03** | **-0,41** |
| **GABA** | **0,02** | **0,42** | 0,69 | -0,13 | 0,10 | -0,38 | 0,41 | 0,18 |
| **Glucose** | 1,00 | 0,15 | 0,34 | 0,67 | **0,01** | **2,43** | **0,03** | **1,91** |
| **Glyceric acid** | 1,00 | -0,01 | 1,00 | 0,08 | **0,03** | **-0,36** | 0,06 | -0,45 |
| **Glycerol-3-phosphate** | **0,02** | **-0,41** | 0,69 | 0,15 | 1,00 | -0,02 | 0,19 | -0,58 |
| **GTP** | **0,03** | **-0,31** | 0,06 | -0,29 | 0,15 | 0,23 | 0,19 | 0,21 |
| **Inositol** | 0,90 | 0,00 | 1,00 | -0,18 | **0,02** | **-0,40** | 0,19 | -0,23 |
| **NADH** | 0,56 | 0,12 | 0,34 | 0,26 | 0,10 | -0,47 | **0,02** | **-0,62** |
| **Phosphoric acid** | 0,29 | -0,03 | 0,11 | -0,47 | **0,02** | **-0,28** | 0,90 | 0,16 |
| **Ribose 5 Phosphate** | 0,90 | 0,16 | 0,69 | 0,00 | **0,02** | **-0,61** | 0,19 | -0,44 |
| **TMA** | 0,11 | -0,19 | 0,06 | 0,19 | **0,01** | **0,46** | 0,73 | 0,08 |

**Table S6: Significant GFP-SYBU interacting proteins in COV318 cells**

| Gene & Synonyms | GFP SYBU/GFP | | | Description |
|---|---|---|---|---|
| | Ratio | Log2 | Adj. p-value | |
| HSPA8 | 10,42840281 | 3,38244631 | 7,28E-157 | Heat shock cognate 71 kDa protein |
| HSPA1B | 5,588902144 | 2,482564915 | 1,17E-95 | RecName: Full=Heat shock 70 kDa protein 1B |
| VDAC1 | 3,531594266 | 1,820319606 | 2,59E-60 | Voltage-dependent anion-selective channel protein 1 |
| HSPA5 | 1,988459132 | 0,991650911 | 2,62E-60 | 78 kDa glucose-regulated protein |
| VDAC3 | 9,473515079 | 3,243899827 | 9,45E-56 | Voltage-dependent anion-selective channel protein 3 |
| VDAC2 | 9,359437348 | 3,226421803 | 4,42E-47 | Voltage-dependent anion-selective channel protein 2 |
| IMMT | 1,901132186 | 0,926858846 | 3,23E-31 | MICOS complex subunit MIC60 |
| BAG2 | 53,3892792 | 5,738478166 | 7,36E-31 | BAG family molecular chaperone regulator 2 |
| HK1 | 4,807415088 | 2,265261376 | 1,09E-27 | Hexokinase-1 |
| MAPRE1 | 4,438655691 | 2,150122802 | 1,09E-27 | Microtubule-associated protein RP/EB family member 1 |
| RUVBL2 | 1,901305481 | 0,926990347 | 4,95E-21 | RuvB-like 2 |
| COPE | 1,944364375 | 0,959298606 | 2,25E-18 | Coatomer subunit epsilon |
| DNAJA3 | 2,113913989 | 1,079916677 | 9,59E-17 | DnaJ homolog subfamily A member 3, mitochondrial |
| PSMC2 | 1,907291375 | 0,93152526 | 3,00E-15 | 26S protease regulatory subunit 7 |
| USP7 | 4,323892607 | 2,112330691 | 4,41E-15 | Ubiquitin carboxyl-terminal hydrolase 7 |
| CSNK2A2 | 2,192447467 | 1,132542275 | 1,31E-12 | Casein kinase II subunit alpha' |
| DYNLL1 | 54,88564838 | 5,778357054 | 3,22E-10 | Dynein light chain 1, cytoplasmic |
| DNAJC13 | 3,341632799 | 1,740553209 | 3,70E-09 | DnaJ homolog subfamily C member 13 |
| CUL1 | 2,809253726 | 1,490186932 | 1,33E-08 | Cullin-1 |
| CSNK2B | 2,936111056 | 1,553906538 | 6,97E-06 | Casein kinase II subunit beta |
| GRWD1 | 2,192111776 | 1,132321363 | 7,49E-06 | Glutamate-rich WD repeat-containing protein 1 |
| MTCH2 | 3,90200723 | 1,964216451 | 9,18E-06 | Mitochondrial carrier homolog 2 |
| FAM91A1 | 2,695940495 | 1,430788653 | 5,88E-04 | Protein FAM91A1 |
| PGAM5 | 1,936184532 | 0,953216458 | 9,63E-04 | Serine/threonine-protein phosphatase PGAM5, mitochondrial |
| SLC25A6 | 2,304769008 | 1,204622166 | 1,05E-02 | ADP/ATP translocase 3 |
| TOMM40 | 1,938952941 | 0,955277789 | 1,32E-02 | Mitochondrial import receptor subunit TOM40 homolog |
| CSNK2A1 | 2,225304164 | 1,154002544 | 1,98E-02 | Casein kinase II subunit alpha |
| MRPL3 | 28,80947226 | 4,848471328 | 2,55E-02 | 39S ribosomal protein L3, mitochondrial |
| THEM6 | 2,383322834 | 1,252974386 | 2,85E-02 | Protein THEM6 |
| SAMM50 | 2,079199585 | 1,056028251 | 3,37E-02 | Sorting and assembly machinery component 50 homolog |
| CDK9 | 3,973743365 | 1,990498702 | 3,38E-02 | Cyclin-dependent kinase 9 |
| BGN | ∞ | ∞ | | Biglycan |
| BLOC1S2 | ∞ | ∞ | | Biogenesis of lysosome-related organelles complex 1 subunit 2 |
| BTRC | ∞ | ∞ | | F-box/WD repeat-containing protein 1A |
| CSNK1D | ∞ | ∞ | | Casein kinase I isoform delta |
| DTNBP1 | ∞ | ∞ | | Dysbindin |
| FBXW11 | ∞ | ∞ | | F-box/WD repeat-containing protein 11 |
| HERC5 | ∞ | ∞ | | E3 ISG15--protein ligase HERC5 |
| HSPA2 | ∞ | ∞ | | Heat shock-related 70 kDa protein 2 |
| HSPA6 | ∞ | ∞ | | Heat shock 70 kDa protein 6 |
| KTI12 | ∞ | ∞ | | Protein KTI12 homolog |
| MAPRE3 | ∞ | ∞ | | Microtubule-associated protein RP/EB family member 3 |
| NIN | ∞ | ∞ | | Ninein |
| OBSL1 | ∞ | ∞ | | Obscurin-like protein 1 |
| Q6ZSR9 | ∞ | ∞ | | Uncharacterized protein FLJ45252 |
| RYR1 | ∞ | ∞ | | Ryanodine receptor 1 |
| SNAPIN | ∞ | ∞ | | SNARE-associated protein Snapin |
| SYBU | ∞ | ∞ | | Syntabulin |
| TRIL | ∞ | ∞ | | TLR4 interactor with leucine rich repeats |
| TRIP12 | ∞ | ∞ | | E3 ubiquitin-protein ligase TRIP12 |
| TRPM6 | ∞ | ∞ | | Transient receptor potential cation channel subfamily M member 6 |
| WDR11 | ∞ | ∞ | | WD repeat-containing protein 11 |
